# Acetylcholine demixes heterogeneous dopamine signals for learning and moving

**DOI:** 10.1101/2024.05.03.592444

**Authors:** Hee Jae Jang, Royall McMahon Ward, Carla E. M. Golden, Christine M. Constantinople

**Affiliations:** Center for Neural Science, New York University; New York, NY 10003

## Abstract

Midbrain dopamine neurons promote both reinforcement learning and movement vigour^1–12^. A major outstanding question is how dopamine-recipient neurons in the striatum parse these heterogeneous signals. Previous work suggests that cholinergic striatal interneurons may play a role, perhaps by gating dopamine-dependent plasticity^13–16^, but this has not been tested in behaving animals. Here we studied rats performing a decision-making task with reward- and movement-related events at distinct time points. Optical measurement of dopamine and acetylcholine release in the dorsomedial striatum (DMS) revealed distinct dynamics at task events. Reward cues evoked cholinergic pauses with different phase relationships relative to dopamine. When dopamine slightly lagged cholinergic dips, dopamine predicted future behaviour, and DMS firing rates on subsequent trials. In contrast, when dopamine slightly preceded cholinergic dips, there was no observable relationship between dopamine and learning. Finally, when dopamine was coincident with cholinergic bursts, it preceded and predicted the vigour of upcoming contralateral orienting movements. Our findings suggest that the precise phase relationship between dopamine and acetylcholine allows dopamine to be used for either movement or learning depending on instantaneous behavioural context.

## Introduction

Midbrain dopamine neurons and their terminals in the striatum are implicated in reinforcement learning and motor control^1–11^. Some evidence suggests that different dopamine neurons may be engaged in learning and movement, such that heterogeneous functions could derive from distinct groups of neurons^4,10,12,17–19^. However, single dopamine neuron axon terminals span millimeters of neuropil^20^, and heterogeneous dopamine signals have been observed at single recording sites in the dorsal striatum^4,10,21^, suggesting that for medium spiny neurons, the principal cells of the striatum, dopamine receptor binding might reflect multiplexed signals for learning and moving. Here we show that acetylcholine release exhibits distinct dynamics when dopamine encodes reward prediction errors (RPEs) or predicts upcoming movement vigour in rats performing a goal-directed decision-making task.

In reinforcement learning, agents learn the value of taking actions in different states; those state-action values are iteratively updated based on RPEs, or the difference between received and expected rewards^22^. A wealth of evidence suggests that dopaminergic neurons instantiate RPEs in the brain^1–8^. One important target for dopamine neurons is the DMS, a striatal region necessary for learning associations between actions and outcomes^23^. Dopamine is thought to drive plasticity at converging synaptic inputs onto medium spiny neurons such that animals will be more (or less) likely to take actions in states that produced positive (or negative) RPEs.

The dopamine system is also implicated in motor control^24^. Degeneration of dopamine neurons projecting to the dorsal striatum is a hallmark of Parkinson’s Disease^25^. Lesions or optogenetic perturbations of dopamine neurons or their axon terminals in the dorsomedial striatum produce gross effects on movement initiation, kinematics, and vigour^9–11^. Studies have found dopamine responses that reflect movements and cannot be easily reconciled with the RPE framework^4,10,21,26^.

Here, we tested the hypothesis that striatal acetylcholine dynamically determines whether dopamine drives learning or movement vigour in rats performing a decision-making task. We developed a task that involves several sequential events on each trial; some events are associated with reward-predictive cues, while other events elicit orienting movements, which require the DMS^27^. We found that dopamine reflected RPEs or predicted the vigour of upcoming movements at distinct time points, and dips in acetylcholine occurred at RPE but not movement events. Dopamine RPEs predicted subsequent behaviour when they slightly lagged cholinergic dips, but not when they slightly preceded cholinergic dips. In contrast, at the onset of contralateral orienting movements, we observed bursts in acetylcholine release. These data suggest that acetylcholine dynamics gate dopamine and striatal function on a moment-by-moment basis.

## Results

### Temporal wagering task provides behavioural readouts of rats’ reward expectations and movement

We trained rats on a self-paced temporal wagering task that manipulated reward expectations by varying the magnitude of offered rewards in blocks of trials^28^ (Fig. 1a,b). Rats initiated trials by poking their nose in the central nose port. This triggered an auditory cue, the frequency of which indicated the volume of water reward offered on that trial (5, 10, 20, 40, or 80 µL). Rats were required to maintain fixation in the central nose poke for ∼1 second while the auditory cue played; leaving the nose port early triggered a white noise sound and terminated the current trial. At the offset of the auditory cue, the reward was assigned randomly to one of the two side ports, indicated by an LED. The rat could wait for an uncued and unpredictable delay to obtain the reward, or at any time could opt out by poking in the other side port to start a new trial. On 15-25% of trials, rewards were withheld to assess how long rats were willing to wait for them (catch trials), providing an analog behavioural measure of how much they valued the reward offer.

**Figure 1:**
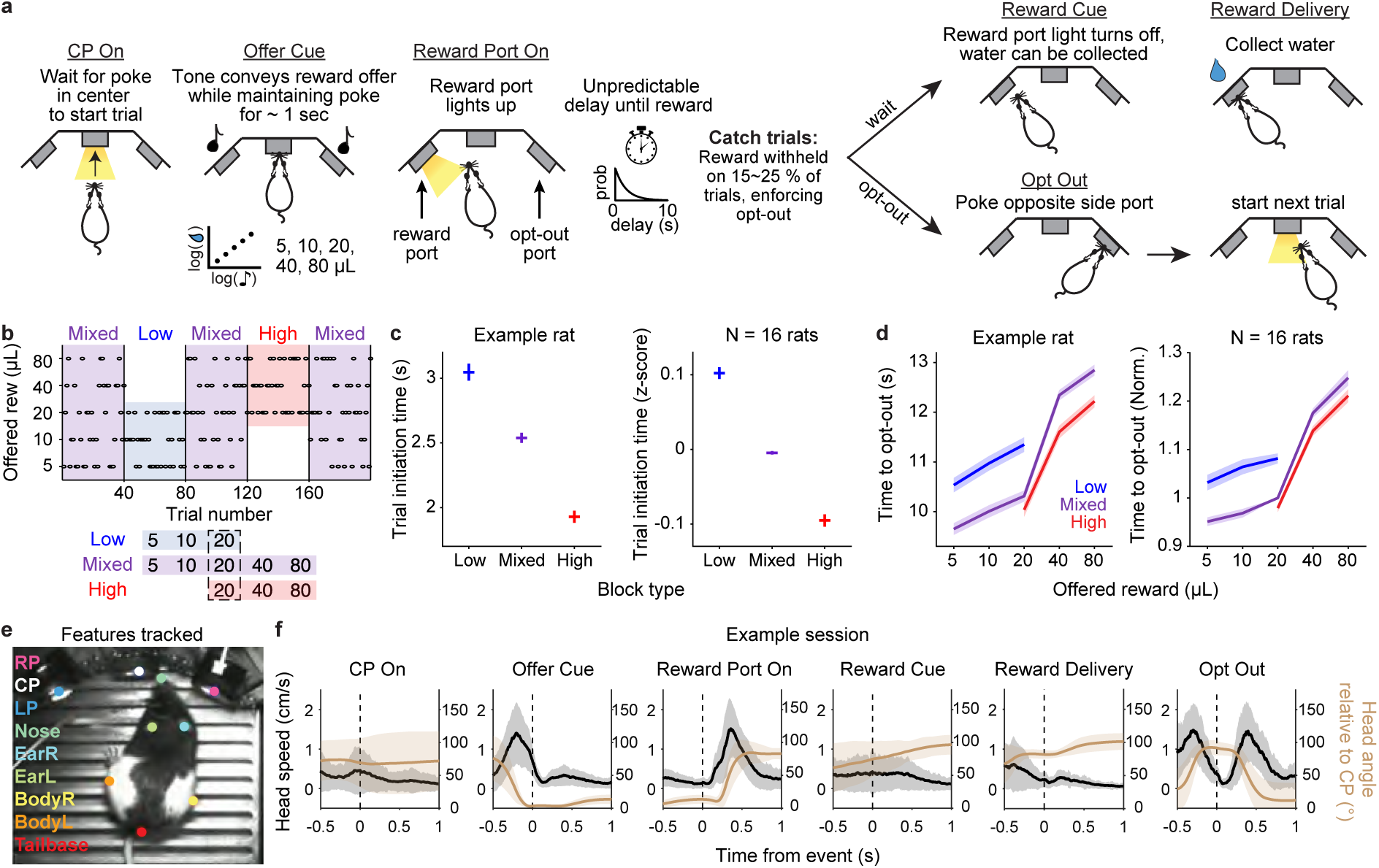
Temporal wagering task provides behavioural readouts of rats’ reward expectations and movement. **a.** Schematic of behavioural paradigm. **b.** Schematic of the block structure of the task. All sessions start with a mixed block and alternate with low or high blocks. Blocks are 40 completed trials. **c.** Left, Average time to initiate trial by block type for an example rat. Trials are pooled across sessions (N = 63 sessions, mean ± s.e.m.). Right, Z-scored trial initiation time by block, averaged across rats (N = 16 rats, mean ± s.e.m.). **d.** Left, Average time to opt-out by offered reward volume in each block for the same example rat in **c**. Trials are pooled across sessions (N = 63 sessions, mean ± s.e.m.). Right, Normalized time to opt-out by offered reward in each block, averaged across rats (N = 16 rats, mean ± s.e.m.). Time to opt-out is divided by the time for 20 µL in mixed block for each rat before averaging. **e.** Features tracked with DeepLabCut using videography. **f.** Head speed (black) and angle relative to the central nose port (brown) aligned to different task events in an example session (mean ± 1 s.d. across trials).

To manipulate their reward expectations, rats experienced uncued blocks of trials in which they were presented with low (5, 10, or 20 µL) or high (20, 40, or 80 µL) volumes of water rewards. These were interleaved with mixed blocks which offered all rewards (Fig. 1b). Rats initiated trials more quickly in high blocks compared to low blocks, providing a behavioural report of their subjective value of the environment (Fig. 1c; N = 16 rats). We also found that rats modulated how long they were willing to wait for rewards based on the average reward in each block (Fig. 1d; N = 16 rats). In previous work, we found that distinct value computations and neural circuits support the decision to initiate a trial versus opt-out during the delay period: initiation times reflect a delta rule and are impacted by perturbing dopamine in the striatum^28–30^, while the decision to opt-out reflects inference of the current block, and relies on the orbitofrontal cortex^28,31^. Given our previous findings that initiation times rely on a dopamine-dependent reinforcement learning algorithm^28–30^, we first focused on that aspect of behaviour.

Deep pose estimation^32^ showed that rats made stereotypical orienting movements at certain task events (Fig. 1e,f). When the side LED lit up indicating the reward port on that trial, rats oriented towards it, and after opting out, they oriented back to the center port to initiate a new trial. Thus, our task includes reward cues that are separable in time from orienting movements, which require the DMS^27^. The task structure enables relating dopamine release to multiple reward-predictive cues and event-aligned movements.

### Dopamine and acetylcholine show distinct dynamics at reward- and movement-associated events

We measured dopamine and acetylcholine release in the DMS in this task. To measure dopamine, we used fibre photometry of an optical dopamine sensor, GRAB*_DA_* (Fig. 2a; N = 6 rats with pAAV-hsyn-GRAB*_DA_*_2_*_h_*, N = 4 rats with pAAV9-hsyn-GRAB*_rDA_*_3_*_m_*). We used a static fluorophore mCherry or isosbestic control to correct for brain motion^33^ in rats injected with DA2h and rDA3m, respectively, with similar results for both methods (Extended Data Fig. 1).

**Figure 2:**
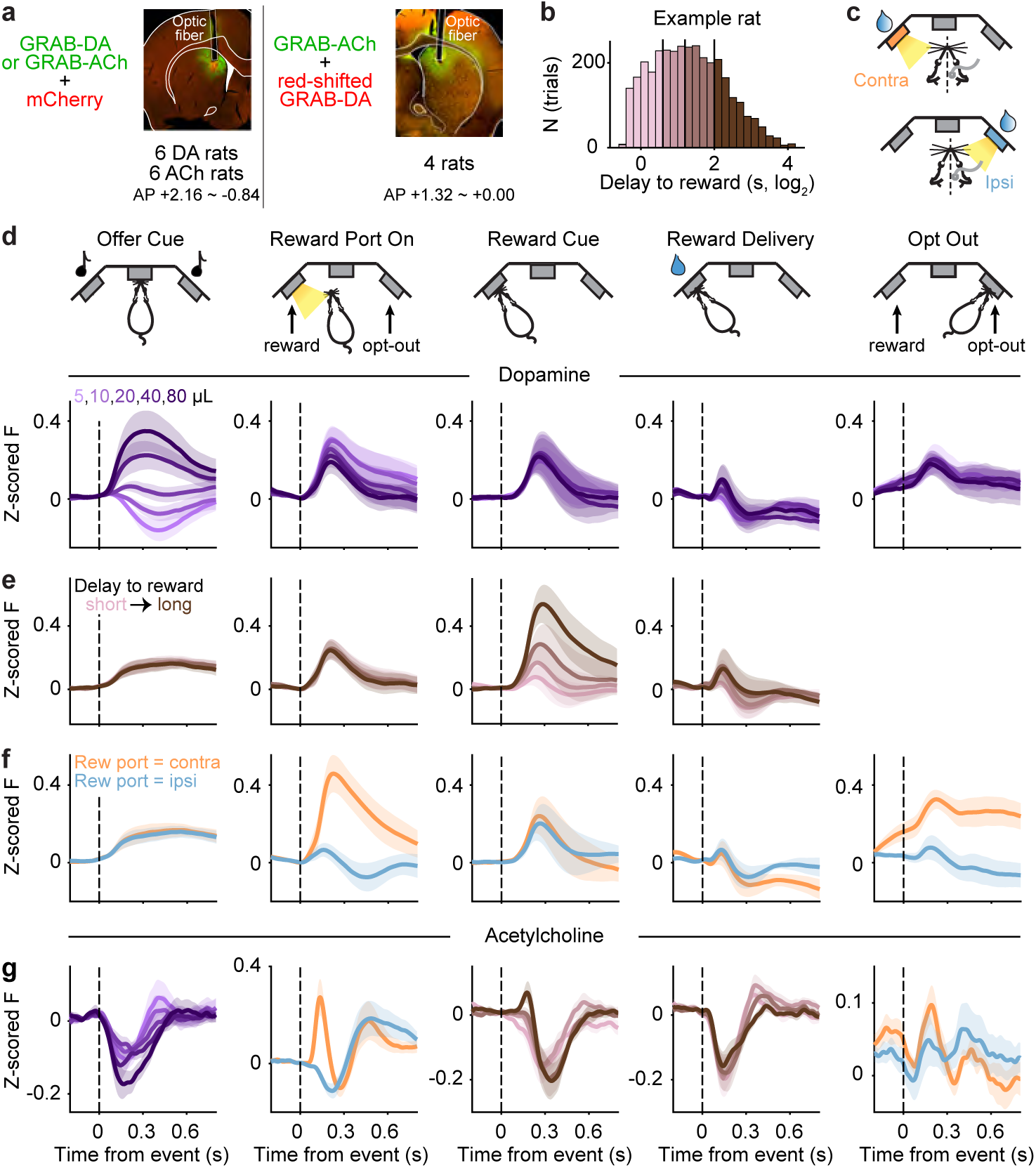
Dopamine and acetylcholine in the DMS show distinct dynamics at reward- and movement-associated events. **a.** Left, Viral strategy to measure dopamine (N = 6 rats, pAAV-hsyn-DA2h) or acetylcholine (N = 6 rats, AAV9-hSyn-ACh4.3) release in the DMS using green fluorescent GRAB sensors and representative histology. Right, Viral strategy to measure dopamine (red, pAAV9-hsyn-rDA3m) and acetylcholine (green, AAV9-hSyn-ACh4.3) simultaneously and representative histology (N = 4 rats). **b.** Distribution of delay to reward, drawn from an exponential distribution, for an example rat. Colours indicate quartiles. **c.** Schematic of contralateral (orange) and ipsilateral (blue) trials shown for a rat implanted in the right hemisphere. The side of implant is counterbalanced across rats. **d.** Average event-aligned dopamine release during mixed blocks for different reward offers (N = 10 rats, mean ± s.e.m.). **e.** Average event-aligned dopamine release split by delay to reward quartiles across all blocks (N = 10 rats, mean ± s.e.m.). **f.** Average event-aligned dopamine release split by contralateral (orange) and ipsilateral (blue) trials (N = 10 rats, mean ± s.e.m.). **g.** Average event-aligned acetylcholine release split by the task variables most strongly encoded by dopamine at each task event (N = 10 rats, mean ± s.e.m.). **d-g.**, Z-scored fluorescence signals are baseline corrected before pooling across rats.

At the offer cue, we observed phasic dopamine release that scaled with the offered reward volume, consistent with an RPE (Fig. 2d). Additionally, because the timing of the rewards was unpredictable and they were omitted on catch trials, we reasoned that when the rat learns reward is available, there should be larger prediction errors for long compared to short delays^34,35^. Binning trials with different reward delays revealed that dopamine release at the reward cue scaled with delay, again consistent with an RPE (Fig. 2b,e). These RPEs were not modulated by the spatial location of the reward port (Extended Data Fig. 2).

However, at other task events when cues elicited stereotypical orienting movements, dopamine release in the DMS showed a larger phasic response for orienting movements that were contralateral to the recording hemisphere^4^ (Fig. 2c,f). This was true when the reward port was assigned and rats oriented toward that port, and following opt-outs, when rats oriented back toward the center port to initiate a new trial. Contralateral orienting signals were strongest in the DMS compared to the ventral and dorsolateral striatum, perhaps reflecting a privileged role for the DMS in executing orienting movements^27^ (Extended Data Fig. 3, 4). Dopamine in the ventral striatum showed similar encoding of RPEs as the DMS^4^ (Extended Data Fig. 3). Dopamine in the dorsolateral striatum, by contrast, did not exhibit RPE encoding at the offer cue (Extended Data Fig. 4). These data are consistent with the DMS representing an intermediate position along the continuum of “reward” to “action” coding in the striatum and its associated dopaminergic innervation^4,10,12,21^. Computing the discriminability index of dopamine responses for different task variables confirmed that DMS dopamine encoded reward volume, delay, and contralateral movements at distinct task events (Methods, Extended Data Fig. 5). Therefore, dopamine in the DMS multiplexed RPEs and contralateral movement signals at distinct time points on single trials.

This raises the question of how dopamine-recipient medium spiny neurons might de-multiplex or demix these heterogeneous signals. To test the hypothesis that acetylcholine may contribute to this process, we next measured acetylcholine release in the DMS using fibre photometry of the GRAB*_ACh_* sensor (Fig. 2a; N = 10 rats injected with AAV9-hSyn-ACh4.3). We found prominent dips in acetylcholine release at cues that elicited dopamine RPEs (Fig. 2g). Notably, the magnitude of the dip scaled modestly with the offered reward volume at the time of the offer cue. In contrast, when the side LED indicated the reward port, we observed a strong burst in acetylcholine release on trials in which that port was contralateral to the recording site (Fig. 2g), when a phasic increase in dopamine release was also observed (Fig. 2f). At opt-out, we also observed a similar but weaker rise in acetylcholine release on contralateral trials (Fig. 2g). The difference in the strength of the acetylcholine burst at the two movement events could reflect more temporal jitter between when rats opt-out and then orient to initiate subsequent trials (Extended Data Fig. 6). Therefore, acetylcholine exhibits dips or bursts when dopamine encodes RPEs versus reflects contralateral movements, respectively. This pattern was observed both when acetylcholine and dopamine were measured in different animals, as well as when we performed dual color imaging of GRAB*_ACh_* and red-shifted GRAB*_DA_* at the same recording site in the DMS (Extended Data Fig 7, 8). This shows that differences in cholinergic and dopaminergic dynamics were not due to slight alterations in the location of the recordings. Recording locations spanned ∼3 mm across the AP axis, so the results appear generalizable across the DMS (see Extended Data Fig 9, 10, 11 for individual histology and dynamics across the AP axis).

### Dopamine RPEs predict learning when is lags cholinergic dips

We next sought to determine whether dopamine at putative RPE events implemented reinforcement learning. Notably, the relative phase relationship between the cholinergic dips and dopamine RPEs was different at the offer cue, when dopamine scaled with reward volume, versus at the reward cue, when dopamine reflected the reward delay. The cross-correlogram between cholinergic dips and dopamine at the offer cue showed that dopamine lagged acetylcholine by ∼100ms at this time point, and the time to the peak (for dopamine) or trough (for acetylcholine) was significantly different (Fig. 3a; N = 10 DA rats, N = 10 ACh rats; two-sided Wilcoxon rank sum: p *<* 0.001). For visualization, we highlight the temporal correlation between large positive RPEs and cholinergic dips, in which dopamine peaks slightly followed cholinergic dips (the temporal relationship was similar for small volumes that produced negative RPEs, but the sign of the correlogram was flipped, Extended Data Fig. 12). At the reward cue, however, the dopamine peak preceded the cholinergic dip by ∼50ms (Fig. 3b; N = 10 DA rats, N = 10 Ach rats; two-sided Wilcoxon rank sum: p *<* 0.001). Examination of the event-aligned responses showed that the dopamine RPE was aligned to the reward cue (when the light turned off after the delay, indicating that reward was baited), but the cholinergic dip was aligned to reward delivery (when the rat poked and triggered the solenoid; Fig. 2e,g). These events occur closely in time, but they are offset and jittered, since the rat has to poke in the nose port following the cue in order to get reward. Therefore, while dopamine appeared to encode an RPE at both the offer and reward cue^29,30^, its relative phase relationship with cholinergic dips was different at these events.

**Figure 3:**
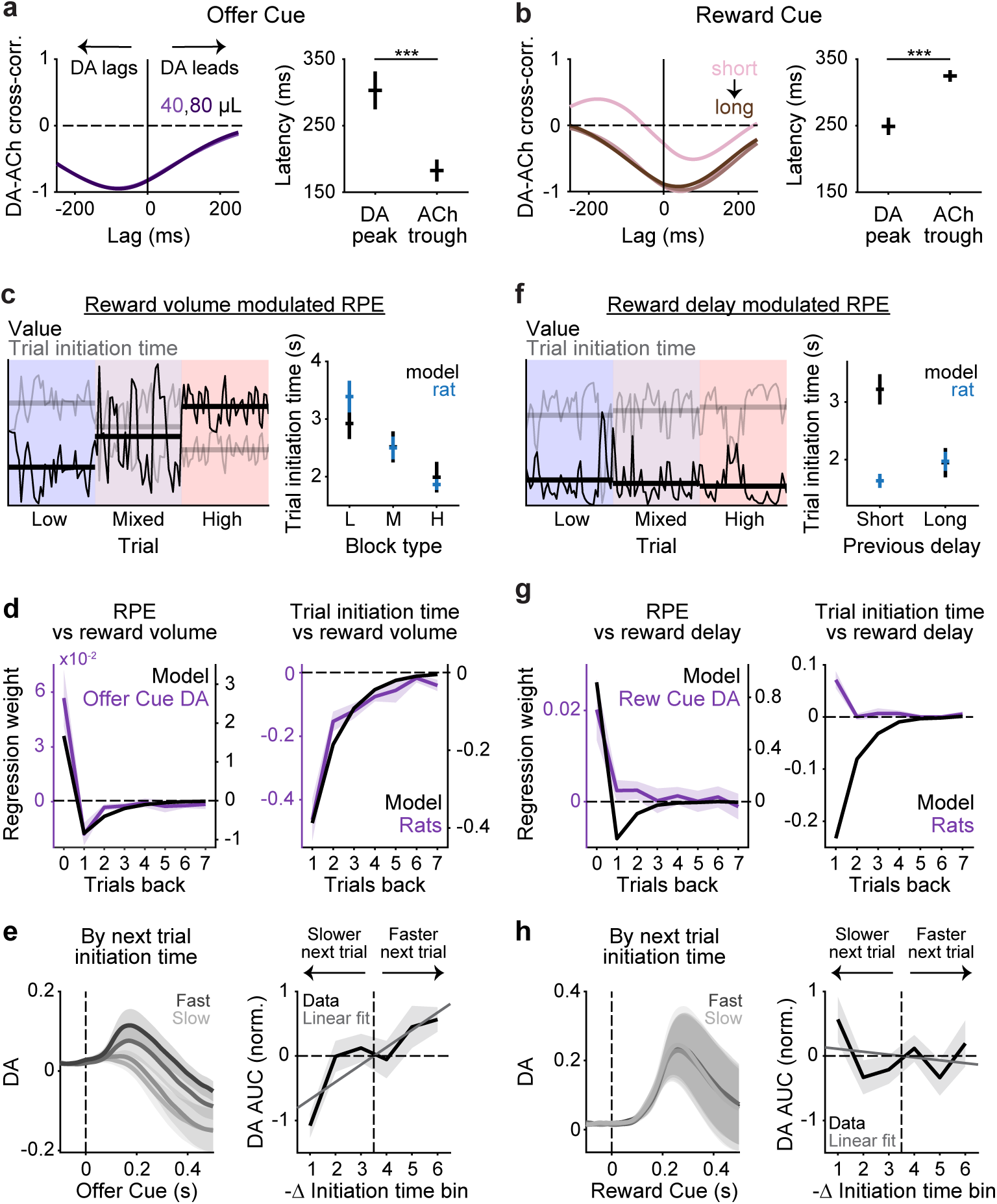
Dopamine correlates with learning when it lags cholinergic dips. **a.** Left, Cross-correlation of rat-averaged dopamine and acetylcholine signals at the offer cue (−0.1-0.5 s) on 40 and 80 µL offer trials in mixed blocks. Negative lag indicates dopamine following acetylcholine signal. Right, Latency to peak/trough in DA/ACh after the onset of the offer cue on 40 and 80 µL offer trials in mixed blocks (N = 10 DA rats, N = 10 ACh rats, mean ± s.e.m.; two-sided Wilcoxon rank sum test: p *<* 0.001). **b.** Left, Same as in **a** but at the reward cue on trials with different reward delay quartiles. Right, Same as in **a** but after the onset of the reward cue on trials with different reward delay quartiles (N = 10 DA rats, N = 10 ACh rats, mean ± s.e.m.; two-sided Wilcoxon rank sum test: p *<* 0.001). **c.** Left, Model predicted value (black) and simulated trial initiation time (grey) for an example rat using reward volume modulated RPE. Solid lines indicate average block values. Right, Trial initiation time by block type averaged across rats (blue) and model simulation using trial statistics from the same rats (black) (N = 10 rats, mean ± s.e.m.). **d.** Left, Dopamine AUC (purple) at the offer cue (0-0.5 s) and model predicted RPE (black) regressed against reward offers in the last 15 trials. Right, Z-scored rat trial initiation time (purple) and model simulation (black) regressed against reward offers in the last 15 trials. Across panels, only mixed blocks are included and 7 trials are shown for visualization purposes (N = 10 rats, mean ± s.e.m.). **e.** Left, Z-scored dopamine at the offer cue conditioned on trial initiation time on the subsequent trial, averaged across rats (N = 10 rats, mean ± s.e.m.). Dark to light colors indicate fastest to slowest quartile of initiation time distribution. Right, Dopamine AUC at the offer cue (0-0.5 s) versus -Δ trial initiation time, averaged across rats (N = 10, mean ± s.e.m., linear mixed-effects model, slope = 0.2690, one-sided p *<* 0.001). **f-h.** Same as in **c-e.** but using reward delay modulated RPE in model simulation. **f,** All blocks are used. **h,** N = 10 rats, mean ± s.e.m., linear mixed-effects model, slope = −0.0442, one-sided p = 0.7385. Across panels, *p *<* 0.05, **p *<* 0.01, ***p *<* 0.001.

Phase relationships varied across subregions of the striatum. In contrast to the DMS, dopamine and acetylcholine in the dorsolateral striatum exhibited an antiphase relationship at the time of reward delivery (Extended Data Fig. 13). This suggests that the phase relationship may vary depending on the behavioural engagement of different parts of the striatum.

We previously found that trial initiation times rely on a dopamine-dependent reinforcement learning algorithm^28–30^, and are causally impacted by dopamine perturbations^30^, so we next sought to relate RPEs at the offer and reward cue to initiation time behaviour. We modeled rats’ trial initiation times as a linearly decreasing function of the value of the environment, which was iteratively updated by the equation *V_t_*_+1_ = *V_t_* + *αδ*, where the *δ* was an RPE. To evaluate the RPEs at offer cue and reward cue separately, we created two models, in which the RPE either reflected the reward volume (*R_t_* − *V_t_*) or the reward delay. Since we observed large RPEs after long delays (Fig. 2e), the delay model RPE scaled with delay (*Delay_t_* − *V_t_*). We simulated the behaviour of both models using parameters that minimized the difference between the average model predictions and rat’s initiation times as a function of block or previous delay (Fig. 3c,f, Methods). The reward volume RPE model could reproduce the rats’ block-dependent initiation times (Fig. 3c). In contrast, because the delay distribution was fixed across blocks, the delay RPE model could not account for changes in initiation times over blocks (Fig. 3f left). The delay RPE model also predicted faster initiation times following long delays, which was not consistent with rats’ behaviour (Fig. 3f right).

The reward volume RPE model further predicted that RPEs should scale with offered rewards, which was observed at the time of the offer cue (Extended Data Fig. 15a). The model also predicted that RPEs should be inversely related to expectations, or the average reward in each block (Extended Data Fig. 15b, *left*). Indeed, the offer cue dopamine response to 20 µL was the largest in low blocks and negative in high blocks, consistent with a volume-dependent RPE (Extended Data Fig. 15b, *right*; N = 10 rats; two-sided Wilcoxon signed-rank: low vs mixed, p = 0.0039; low vs high: p = 0.0020; mixed vs high: p = 0.0273). The model predicts that RPEs and behaviour will be influenced by previous rewards, with the most recent rewards having the greatest influence and decaying exponentially further in the past (Fig. 3d). Regressing dopamine area under the curve (AUC) and initiation times at the time of the offer cue against previous rewards revealed negative, exponentially decaying influences of previous rewards on phasic dopamine responses (Fig. 3d left), consistent with an RPE, as well as rats’ initiation times (Fig. 3d right). Finally, there was a positive relationship between the magnitude of the phasic dopamine signal at the offer cue and the change in initiation times on the subsequent trial (Methods, Fig. 3e; N = 10 rats; linear mixed-effects model, slope = 0.2690, one-sided p *<* 0.001).

In contrast, dopamine at the reward cue reflected the current reward delay, but not past reward delays, as would be expected from an iterative reinforcement learning algorithm (Fig. 3g left). If large RPEs at the offer cue were increasing the value of the environment, we would expect rats to exhibit faster initiation times following long delays (i.e., regressing initiation time against delay should yield negative coefficients). However, the previous delay had a small positive coefficient (Fig. 3g right). Finally, there was no relationship between the magnitude of the phasic dopamine signal at the reward cue and the change in initiation times on the subsequent trial (Methods, Fig. 3h; N = 10; linear mixed-effects model, slope = −0.0442, one-sided p = 0.7385).

Collectively, these results show that when dopamine slightly lags cholinergic dips at the offer cue, it predicts subsequent initiation times, consistent with acting as an RPE that updates the value of the environment. In contrast, when RPEs slightly precede cholinergic dips, we did not find evidence that it acted as an RPE to update the value of the environment for initiation times.

### Dopamine fails to predict learning when it precedes cholinergic dips

We next sought to determine what might be learned from the reward cue RPE, which scaled with the reward delay. Because we did not observe a strong relationship between reward delay and trial initiation times, we next examined how long rats were willing to wait before opting out of the trial. We have previously modeled this aspect of behaviour with a time-varying function that reflects the value of waiting^28^. The starting point reflects the reward offer, and the shape reflects the probability of reward over time, dictated by the catch probability, *p*, and the time constant of the exponential delay distribution, *τ* (Fig. 4a). The model opts-out when the value of waiting falls below the value of the environment, implemented as one of three block-specific thresholds (Fig. 4b). The model captures various aspects of opt-out decisions, including sensitivity to blocks and reward offers (Fig. 1d), and dynamics of behavioural changes at various block transitions^28,31^. Manipulating the catch probability *p* changed how long rats were willing to wait (Fig. 4c; N = 53 rats; two-sided Wilcoxon signed-rank: p *<* 0.001 for all reward offers), consistent with the model^28^ (Fig. 4d), suggesting that rats learned this time-varying function through experience. We thus considered the possibility that the delay RPE might update rats’ estimates of the value of waiting, i.e., temporal reward expectations during the delay. According to temporal difference learning (and belief-state variants^35^), large RPEs after a long delay should increase the value of the states preceding the reward, which in our task should cause rats to wait longer on the subsequent trial (Fig. 4e). However, there was no difference in how long rats waited for rewards in mixed blocks, conditioned on whether the reward delay on the previous trial was in the top versus bottom delay quartile (Fig. 4f). Regressing the time to opt-out against past reward delays yielded negligible coefficients (Fig. 4g). In contrast, how long rats waited in mixed blocks before opting-out showed a strong relationship with the current reward offer (Fig. 4g; N = 16 rats; t-test: p *<* 0.001).

**Figure 4:**
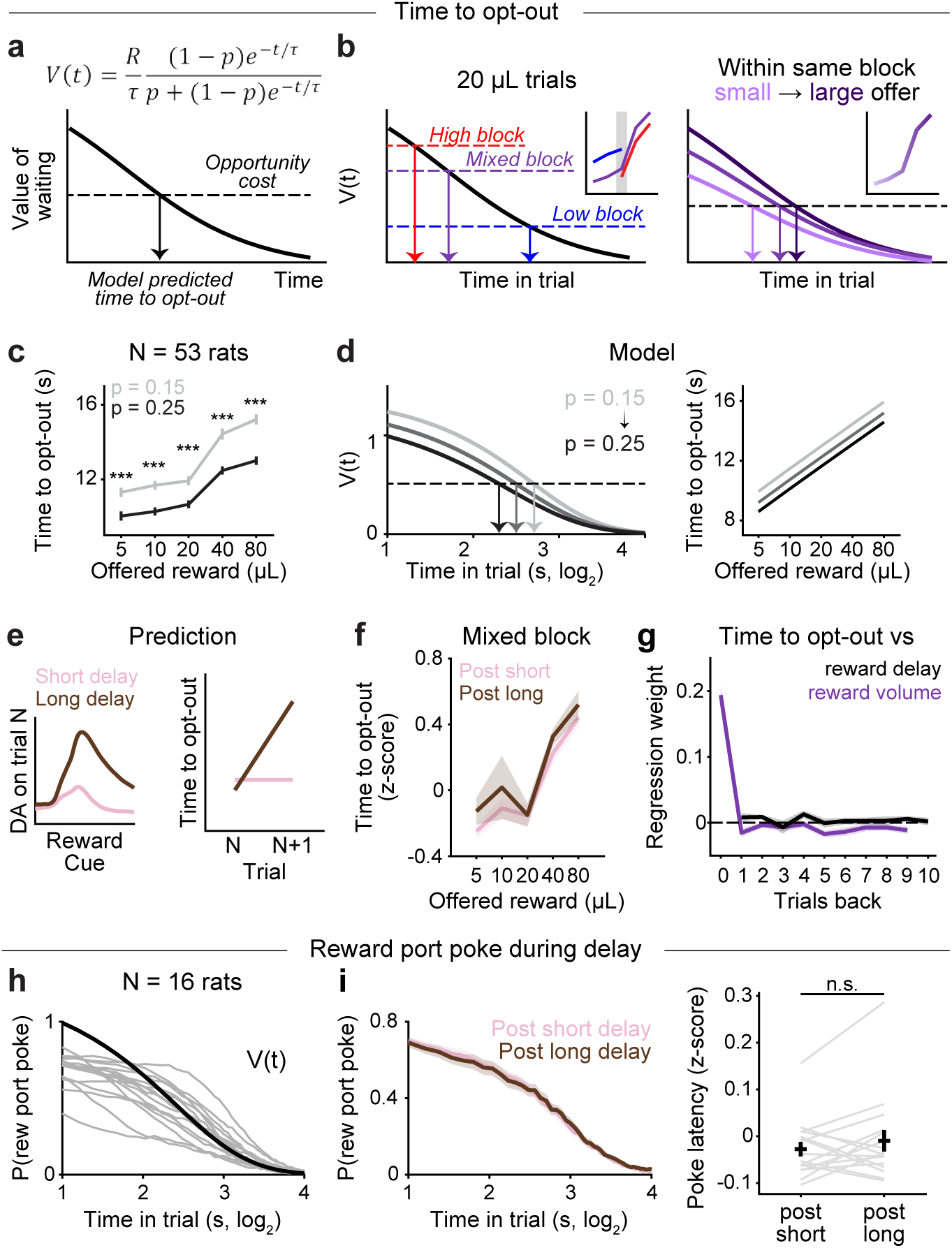
Dopamine fails to predict learning at reward cue when it precedes cholinergic dips. **a.** Value of waiting during the delay period on a given trial as a function of *p*(*catch*) (probability of the trial being unrewarded), and *τ* (time constant of the exponential delay distribution). The model opts out when the value of waiting falls below the opportunity cost, which is fixed for a given block. **b.** Left, Block-specific opportunity cost predicts different time to opt-out on 20 µL offer trials, consistent with rats’ behaviour. Right, In a given block, value function predicts monotonic relationship between reward offer and time to opt-out, consistent with rats’ behaviour. Insets show rats’ behaviour from Fig. 1d. **c.** Average time to opt-out in mixed blocks for rats tested with catch probabilities of 0.15 (light gray) and 0.25 (black). N = 53 rats, mean ± s.e.m., two-sided Wilcoxon signed-rank: p *<* 0.001 for all reward offers. **d.** Value function when *p*(*catch*) changes (left) and model-predicted time to opt-out in mixed blocks for different values of *p*(*catch*) (right). **e.** Large RPEs after long delay trials (brown) should increase the value of preceding states and thus the time to opt-out on the subsequent trial. Negligible DA evoked on short delay trials (pink) predicts little or no change in time to opt-out on the subsequent trial. **f.** Z-scored time to opt-out during mixed blocks conditioned on reward delay on the previous trial, averaged across rats (N = 16 rats, mean ± s.e.m.). Short (pink) and long (brown) delays are defined as the bottom/top quartile of the reward delay distribution across sessions for each rat. **g.** Time to opt-out regressed against reward delay (black) and reward volume (purple) in the past 15 trials during mixed blocks, averaged across rats (N = 16 rats, mean ± s.e.m., t-test: time to opt-out vs reward delay, trial back = 2, p = 0.0223; trial back = 4, p = 0.0398; time to opt-out vs reward volume, trial back = 0, p *<* 0.001; trial back = 1, p = 0.0344; trial back = 5, p = 0.0046; trial back = 6, p = 0.0473). Only ten trials are shown for visualization purposes. **h.** Probability of nose poking in the reward port during the delay period in mixed blocks overlaid with task-defined value function (*τ* = 2.5, *p*(*catch*) = 0.25). Grey lines show individual rats (N = 16 rats). **i.** Left, Probability of nose poking in the reward port during the delay period following trials in the bottom (pink) and top (brown) quartile of reward delay distribution (N = 16 rats, mean ± s.e.m.). Right, Z-scored latency to poke in the reward port after the reward port light turns on following short vs long reward delay trials, averaged across rats (N = 16 rats, mean ± s.e.m., two-sided Wilcoxon signed-rank: p = 0.3011).

We also measured the probability that rats were poking their nose in the reward port in different time bins during the delay period. This probability exponentially decayed over time in line with the veridical reward hazard rate, defined by *p* and *τ* (Fig. 4h). Notably, there was no difference in the poke probability following reward delays in the top versus bottom delay quartiles (Fig. 4i left). We also compared the latency to initially poke in the reward port, the vigour of which could be influenced by temporal reward expectations. Again, there was no difference in initial poke latencies following long or short reward delays (Fig. 4i right; N = 16 rats; two-sided Wilcoxon signed-rank test: p = 0.3011).

Therefore, when dopamine encoded a delay-dependent RPE that slightly preceded cholinergic dips at the reward cue, we failed to find a relationship between dopamine and behaviour on subsequent trials, including rats’ trial initiation times, time to opt-out, probability of poking in the reward port during the delay, or initial latency to poke in the reward port. While it is possible that this RPE promoted some type of learning that we could not resolve, this negative result is in contrast with dynamics at the offer cue; when dopamine slightly lagged cholinergic dips at that timepoint, it predicted trial-by-trial changes in initiation times. These data show that the precise phase relationship between cholinergic dips and dopamine RPEs gates whether those RPEs actually promote learning.

### Dopamine promotes plasticity when coincident with cholinergic dips

Since phasic dopamine at the time of the offer cue is consistent with an RPE, we hypothesized that it might produce plasticity in the recipient DMS cells^36–38^. Dopamine receptor binding can influence the excitability of medium spiny neurons^39,40^, or modify the strength of convergent synaptic inputs^36–38^. However, it has remained unclear how and when RPE-like updating of DMS activity actually occurs in vivo during behaviour. For RPE updating to be useful, it must be rapidly occurring but also enduring, and it is not clear how the timescales of conventional short-term and long-term plasticity allow behaviourally useful changes to occur in this manner.

We recorded single unit action potentials using Neuropixels probes implanted in the DMS (Fig. 5b; N = 8 rats). Geometric constraints for chronic implants precluded simultaneous photometry and Neuropixels recordings at the same site. Therefore, we used the trial-by-trial change in initiation times as a behavioural proxy for RPEs on each trial (Fig. 3e). We observed trial-by-trial changes in firing rate following increases or decreases in initiation times at the time of the offer cue, but not at subsequent events (Fig. 5c). To quantify this effect, we regressed trial-by-trial changes in average firing rates for 300 ms after the onset of each event against putative RPEs (-Δ initiation times; Methods). The highest proportion of cells had significant slope parameters at the time of the offer cue (51.6%, 130/252 units) with 53.1% (69/130 units) having a significantly positive slope (e.g., example cell in Fig. 5d) and 46.9% (61/130 units) having a significantly negative slope (Fig. 5e,f). It is possible that these groups reflect neurons expressing D1- and D2-receptors, respectively. The relationship between putative RPEs and trial-by-trial changes in firing rate decreased at events that were further from the offer cue, suggesting that plasticity was temporally specific to the time of the RPE. We confirmed that the results in Fig. 5c-f reflect RPEs from the previous trial rather than the initiation time on the current trial (Extended Data Fig. 16).

**Figure 5:**
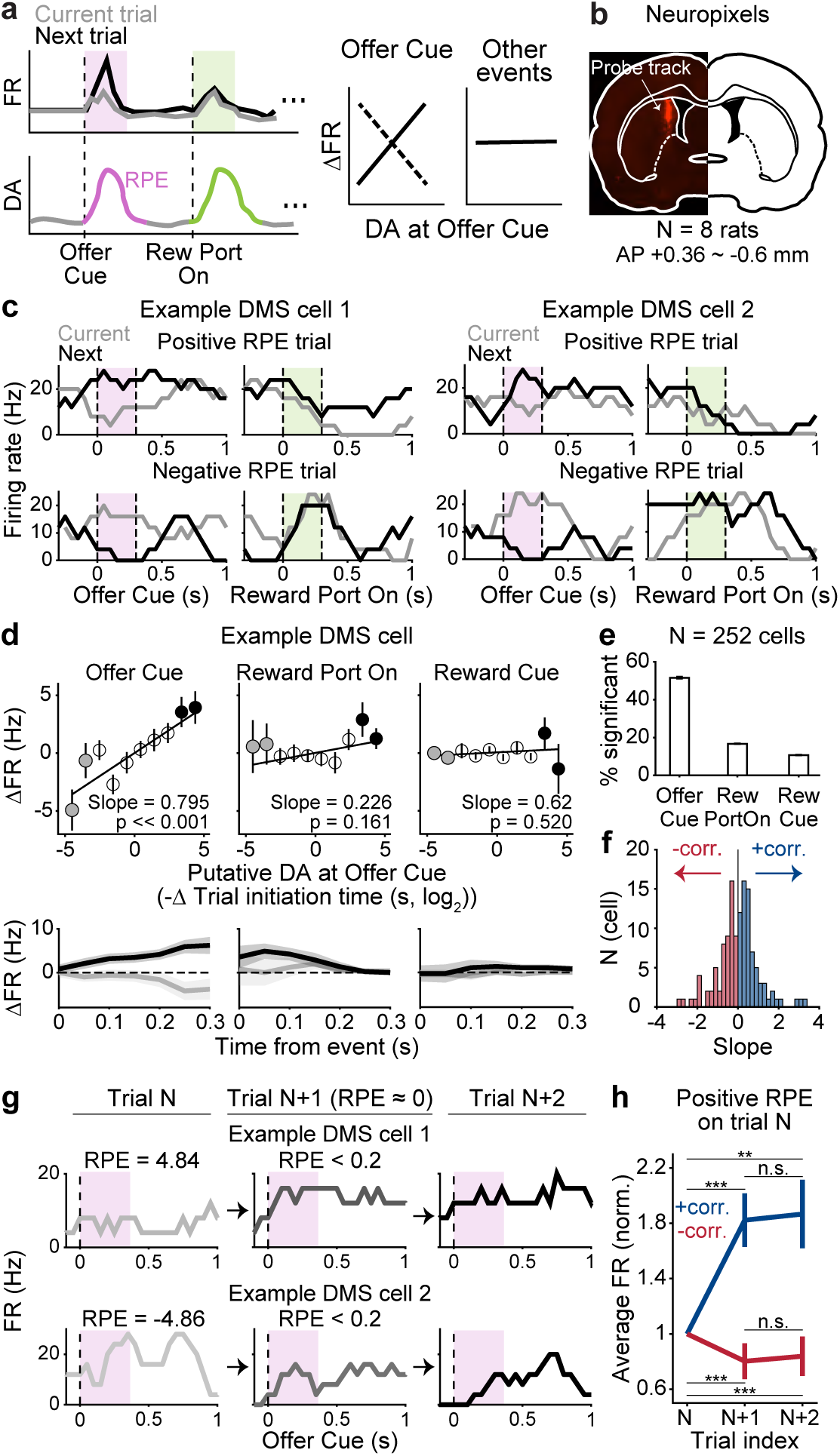
Behavioural proxy for dopamine predicts DMS firing rate on subsequent trials. **a.** Schematic of the prediction for state- or input-specific dopamine-dependent synaptic plasticity where RPE at the time of the offer cue produces trial-by-trial changes in firing rates at that specific event but not at other states. **b.** Example histology of Neuropixels probe track in the DMS. **c.** Firing rate of two example cells on current (grey) and the subsequent trial (black) following a positive and negative RPE. Shaded boxes indicate the time window used to compute average firing rate (0-0.3 s) in the following panels. **d.** Top, Trial-by-trial changes in average firing rate surrounding each task event (0-0.3 s) versus putative dopamine at offer cue (-Δ*log*_2_ trial initiation time) for an example cell. Solid lines show the best fit line (t-test on slope: Offer Cue, p *<* 0.001; Reward Port On, p = 0.161; Reward Cue,p = 0.520) and the filled black/grey circles indicate trials with large positive/negative RPEs, respectively. Trials are binned in 15 linearly spaced intervals for visualization (mean ± s.e.m.). Bottom, Change in average firing rate on trials with large positive (black) and negative RPEs (grey) for the same example cell in the top panel (mean ± s.e.m. across trials). **e.** Fraction of cells with significant slopes at different task events. Error bars are binomial confidence intervals. **f.** Histogram of slopes for significantly modulated cells at the offer cue (red: positively modulated by RPE, blue: negatively modulated by RPE). **g.** Firing rate of two example cells aligned to the onset of offer cue on 3 consecutive trials where a big positive/negative RPE on the first trial is followed by a negligible RPE on the second trial. **h.** Change in average z-scored firing rate at offer cue (0-0.3 s) of positively (blue) and negatively (red) modulated cells on 3 consecutive trials for which a large positive RPE (top 15 %) on trial N is followed by a negligible RPE on trial N+1 (N = 145/164 trial sequences for +corr./-corr., mean ± s.e.m., two-sided Wilcoxon signed-rank: positively modulated cells, trial N vs N+1, p = 0.0004; trial N vs N+2, p = 0.0065; trial N+1 vs N+2, p = 0.4169; negatively modulated cells, trial N vs N+1, p *<* 0.001; trial N vs N+2, p *<* 0.001; trial N+1 vs N+2, p = 0.1898). Across panels, *p *<* 0.05, **p *<* 0.01, ***p *<* 0.001.

To characterize the timescale of these effects, we identified sequences of trials in which a large positive/negative RPE (top/bottom 15%) on trial N was followed by a negligible RPE (|RPE| *<* 0.2) on trial N+1. If dopamine effects are long-lasting, the change in firing rate from trial N to trial N+1 should persist to trial N+2, since there was a negligible RPE on the intervening trial. Fig. 5g shows the firing rates of two example neurons on such a trial sequence. We analyzed firing rates for all such sequences separately for cells with positive and negative firing rate correlations with RPEs. In both groups, positive RPEs on trial N changed firing rates on trial N+1, and that change persisted to trial N+2 (Fig. 5h, two-sided Wilcoxon signed-rank: positively modulated cells, trial N vs N+1, p = 0.0004; trial N vs N+2, p = 0.0065; trial N+1 vs N+2, p = 0.4169; negatively modulated cells, trial N vs N+1, p *<* 0.001; trial N vs N+2, p *<* 0.001; trial N+1 vs N+2, p = 0.1898). These data are consistent with RPEs producing lasting changes in task-evoked firing rates that endured over trials. This result generally held across a range of thresholds used to select RPEs on trial N (Extended Data Fig. 17a). Negative RPEs produced more variable effects on trial-by-trial spiking for these specific sequences of trials (Extended Data Fig. 17b), possibly reflecting different reliability of plasticity mechanisms following increases versus decreases of dopamine. Nonetheless, positive RPEs in the DMS modulate evoked firing on subsequent trials at the time of the offer cue but not at neighbouring task events, including the reward cue (Fig. 5c-e), and those changes persist over trials (Fig. 5g,h).

### Dopamine predicts movement vigour when coincident with cholinergic bursts

Next, we sought to characterize the contralateral selective dopamine signal at events accompanying orienting movements, when dopamine was coincident with cholinergic bursts (Fig. 6a,b). The time to peak of the dopamine signal at these events preceded that in head speed by ∼100 ms (Fig. 6c,e; N = 24 sessions from 5 rats; t-test: Reward Port On, p = 0.0135; Opt Out, p *<* 0.001). Furthermore, the amplitude of the dopamine signal at these events predicted the vigour of the subsequent orienting movement. When the reward port was assigned by the LED, rats quickly oriented towards that port. The AUC of the dopamine signal at this event was significantly larger when reaction times were in the fastest compared to the slowest quartile (Fig. 6d; N = 10 rats; two-sided Wilcoxon signed-rank: black dots indicate time points with p *<* 0.05). This was true when we restricted trials to those with the same reward offer volume (Extended Data Fig. 20b; N = 10 rats; two-sided Wilcoxon signed-rank: p = 0.0137). Notably, neither movement vigour (Extended Data Fig. 19) or dopamine release (Extended Data Fig. 20a) at this timepoint were positively correlated with the reward offer volume, consistent with this orienting movement reflecting a prepotent response to a salient stimulus^41^. Therefore, variability in movement speed and dopamine release were independent of reward expectations at this event. Similarly, after rats opt-out, they orient back to the central nose port to start a new trial, and the AUC of the dopamine signal at opt-out was also larger when rats were faster to initiate the next trial (Fig. 6f; N = 10 rats; two-sided Wilcoxon signed-rank: black dots indicate time points with p *<* 0.05). Therefore, the phasic dopamine signal at these movement-associated events preceded and predicted the vigour of upcoming contralateral orienting movements.

**Figure 6:**
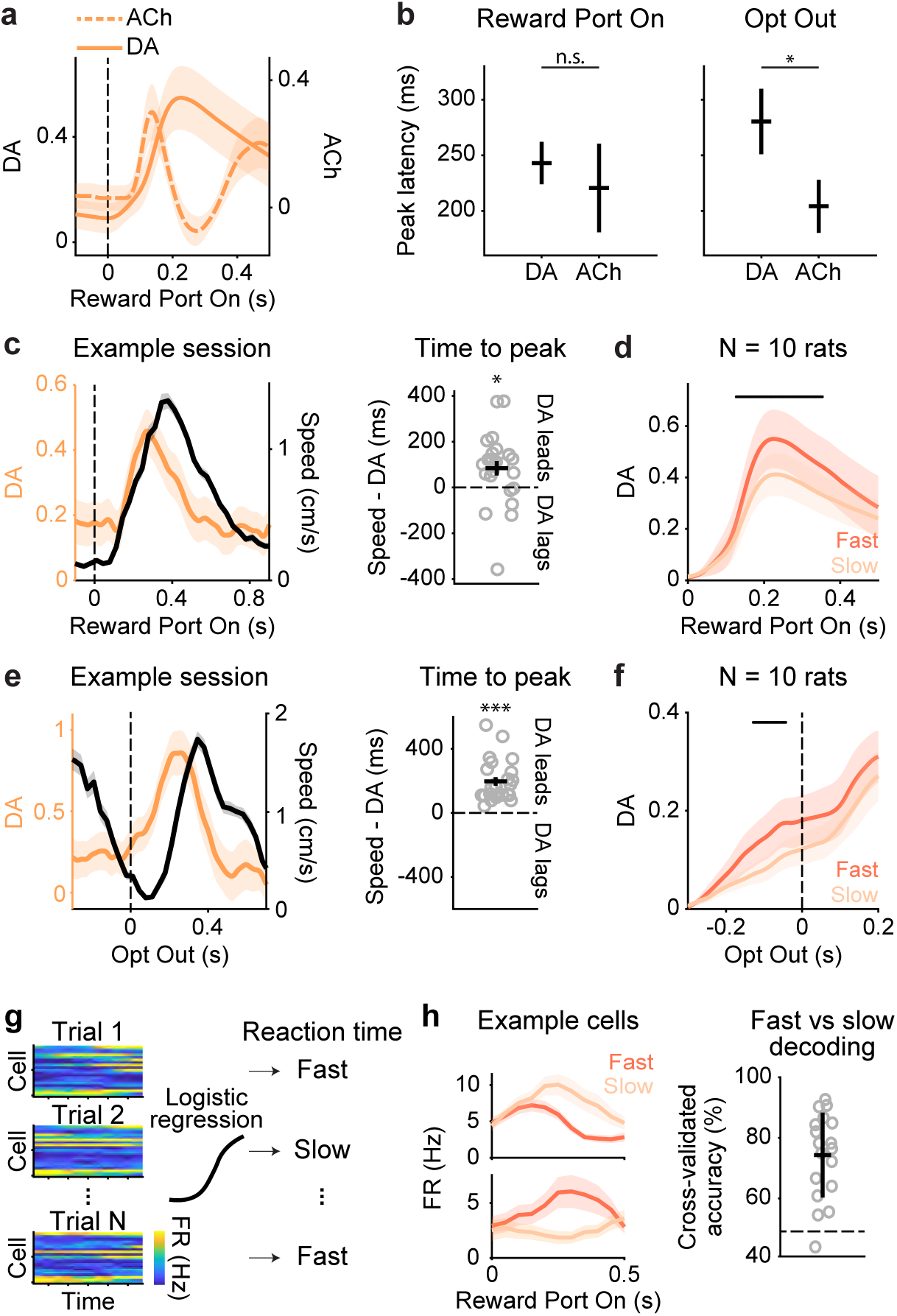
Dopamine predicts movement vigour when coincident with cholinergic bursts. **a.** Z-scored DA (solid orange) and ACh (dashed orange) responses on contralateral trials aligned to when the side LED turns on, averaged across rats (N = 10 DA rats, N = 10 ACh rats, mean ± s.e.m.). **b.** Latency to peak in dopamine and acetylcholine after the side LED turns on (left) and when rats opt out (right) on contralateral trials, averaged across rats (N = 10 DA rats, N = 10 ACh rats, mean ± s.e.m.; two-sided Wilcoxon rank sum test: Reward Port On, p = 0.1109; Opt Out, p =0.0483). **c.** Left, Dopamine release on contralateral trials (orange) when the side LED turns on, overlaid with head speed (black) in an example session (mean ± s.e.m. across trials). Right, Difference in time to peak in dopamine compared to head speed (N = 24 sessions from 5 rats, mean ± s.e.m. across sessions, t-test: p = 0.0135). Empty circles indicate individual sessions. **d.** Z-scored dopamine release on contralateral trials aligned to side LED onset, split by reaction time (fast: bottom quartile; slow: top quartile). Reaction time quartiles were defined within each session, and session-averaged responses were computed for each rat before pooling across rats (N = 10 rats, mean ± s.e.m.). Black line indicates time points with significant differences between fast versus slow trials (Two-sided Wilcoxon signed-rank: p *<* 0.05). **e.** Same as in **c** but for opt-out (t-test: p *<* 0.001). **f.** Same as in **d** but at opt-out split by the speed of the following trial initiation (fast: bottom quartile; slow: top quartile; N = 10 rats, mean ± s.e.m.). Sessions were aggregated to define trial initiation quartiles for each rat since number of opt-out trials are limited in each session. **g.** Schematic of a logistic regression decoder which takes population responses 0-0.5 s at the onset of side LED as input to predict reaction time on a given contralateral trial. **h.** Left, Average firing rate of two simultaneously recorded example cells on fast (dark orange) versus slow (light orange) reaction time trials (mean ± s.e.m. across trials). Right, Average cross-validated classification accuracy of a logistic regression decoder. Dashed horizontal line indicates chance performance (N = 18 sessions from 8 rats, mean ± s.e.m. across sessions). Grey circles indicate individual sessions. Across panels, *p *<* 0.05, **p *<* 0.01, ***p *<* 0.001.

To test whether dopamine at this event promoted plasticity, we tested for a significant correlation between trial-by-trial changes in the firing rates of DMS cells and dopamine release (using the reaction time on the previous contralateral trial as a proxy for dopamine). In contrast to RPE events, when behavioural proxies for dopamine predicted trial-by-trial firing rates, we found that for the overwhelming majority of cells, there was no significant correlation (Extended Data Fig. 18). However, we found that DMS cells encoded movement vigour at this timepoint. We trained a logistic regression decoder to predict whether the reaction time on each trial was in the fastest or slowest quartile, based on the firing rates of simultaneously recorded neurons (Methods, Fig. 6g). Cross-validated classification accuracy was high and above chance in 17 of 18 sessions (Fig. 6h), suggesting that dopamine and DMS spiking at movement-associated events predicted the vigour of upcoming contralateral orienting movements but did not result in persistent changes in task-evoked spiking of DMS cells.

## Discussion

Although cholinergic interneurons only comprise 1-3% of neurons in the striatum^42^, they are thought to be critical for basal ganglia function. Tonically active neurons (putative cholinergic interneurons) develop pauses in firing following cues that predict appetitive or aversive outcomes^15,43–48^, suggesting that cholinergic pauses are important for or reflect learning^49,50^, possibly through local modulation of dopamine release^51–53^. While some studies reported weak relationships between cholinergic firing and movement^15,43,44,47^, others have found signals that reflect motor responses^48,54,55^. Our data suggest that the relative phase relationship between dopamine and acetylcholine^56–58^ is key for determining dopamine’s action on recipient neurons. Recent work suggests that temporal dynamics of dopamine release can vary depending on the behavioural engagement of striatal subregions^59^. We speculate that modulation of the phase relationship between dopamine and acetylcholine provides a circuit mechanism for arbitration between behavioural controllers originating from different striatal subregions; when striatal subregions do not support behaviour, or dopamine-dependent plasticity in those subregions is not desirable, dopamine RPEs may lead (instead of lag) cholinergic dips. Moreover, dopamine signals that seem inconsistent with RPE (for instance, phasic bursts to cues that predict aversive outcomes^60^) may be more interpretable in the context of concurrent cholinergic dynamics.

There is a growing appreciation that dopamine neurons exhibit heterogeneous responses during complex tasks^10,12^, and in some cases different response profiles might map onto transcriptionally defined dopamine cell types^61–63^. Our findings show that in the dorsomedial striatum, dopamine RPE and movement-predictive signals are present at the same recording site. While these signals might reflect inputs from different dopamine neurons, if axonal arbors are overlapping, which is likely^20^, then medium spiny neurons will receive heterogeneous dopamine inputs. The temporal relationship between dopamine and acetylcholine release that is important for medium spiny neurons likely reflects several complex factors, including receptor affinity, efficacy and dynamics of reuptake, and the timescale of interactions between second messengers downstream of dopaminergic and cholinergic GPCRs. Here, we showed that dopamine predicted ongoing movement vigour or subsequent behaviour depending on its phase relationship with acetylcholine, but the ground truth temporal dynamics experienced by medium spiny neurons at these events may be different than what we measured. For instance, indicator kinetics presumably impacted the timing of the signals we measured. We used both high affinity (GRAB*_DA_*_2_*_h_*) and medium affinity sensors (GRAB*_rDA_*_3_*_m_*) to measure dopamine, which have been reported to have similar onset kinetics^64^. Indeed, there were no significant differences in timing of positive dopamine RPEs with the medium versus high affinity sensors (Extended Data Fig. 14), although offset kinetics might be more sensitive to sensor affinity^64^.

While it is unclear why the reward cue RPE did not impact behaviour, at least by the metrics we evaluated, we speculate that it may be because knowing the reward delay on trial *t* does not help one predict the reward delay on trial *t* + 1; the delay is inherently stochastic, and its uncertainty is irreducible (i.e., it reflects expected uncertainty^29,65^). Therefore, learning from this RPE will not help one maximize future rewards. An interesting future question is how animals might learn that RPEs at different events reflect distinct types of uncertainty^65^, and how cholinergic dynamics evolve over that learning process.

Classic studies suggested that corticostriatal plasticity is governed by a three-factor rule requiring coincident pre- and post-synaptic spiking and dopamine receptor binding^38,66,67^. Recent work has revised this view, arguing that a fourth factor - pauses in cholinergic interneurons – is required for induction of synaptic plasticity^13^. Our findings support this hypothesis, although the mechanism is unclear. We hypothesize that G-proteins downstream of dopamine and acetylcholine receptors have opposing effects on intracellular second messengers such as adenylyl cyclase. For instance, M4 receptors are found on D1-receptor-expressing medium spiny projection neurons^68^. M4 receptors are coupled to G*_i/o_* proteins, and when activated, inhibit adenylyl cyclase and downregulate cAMP. Consequently, M4 receptor activation can counteract the positive cAMP signaling induced by dopamine binding to D1 receptors. This suggests that at tonic concentrations of dopamine and acetylcholine, (short or long-term) potentiation and depression mechanisms are in competition, with phasic dopamine favoring potentiation via D1-mediated PKA activation, and acetylcholine favoring depression through M4 activation. However, cholinergic pauses may relieve M4-mediated suppression of adenylyl cyclase, and allow D1-mediated enhancement of PKA. The mechanism could also be presynaptic: corticostriatal and thalamostriatal afferents express M2 receptors, which suppress glutamatergic transmission, and blocking those receptors promotes LTP^69–71^. Therefore, cholinergic pauses may relieve M2-mediated suppression of glutamatergic inputs to the striatum^69^. The relative phase relationships that gate or block plasticity might reflect the impact of receptors with different affinities (e.g., D1 is low affinity, M2/M4 are high affinity) on the relative timing of intracellular second messengers.

Despite the prominence of the dopamine RPE hypothesis, it has remained unclear how and when RPE-like updating of striatal activity actually occurs during behaviour. While reinforcement learning models of behaviour assume that learning is reflected in trial-by-trial changes in state-action values (Q-values), represented in corticostriatal synaptic weights^72^, studies in slices typically evaluate effects of long-term plasticity protocols on the order of minutes after tetanic stimulation or pre/post spike pairing. Our data suggest that induction of temporally-specific plasticity (either short or long-term) can occur within the seconds-scale inter-trial intervals in our task, while expression of plasticity via changes in striatal firing rates is persistent over trials. The temporal specificity of effects at RPE but not movement events further suggests that trial-by-trial changes in firing rates did not simply reflect dopaminergic modulation of excitability^39,40^. These results add to recent findings in other brain regions of rapid *in vivo* plasticity that operates on behavioural timescales^73^, and validate key assumptions of reinforcement learning accounts of behaviour.

## Methods

### Rats

A total of 82 Long-Evans rats (ages 6–24 months) were used in this study. Of these, 25 rats were used for fibre photometry experiments, 8 for electrophysiology experiments, and 53 for behavioural analysis in Fig. 4c (including 4 rats that also underwent fibre photometry).

Of the 25 rats used for fibre photometry, 16 rats were implanted in the DMS (6 with GRAB*DA*2*h*, 6 with GRAB*ACh*, and 4 with a combination of GRAB*rDA*3*m* and GRAB*ACh*), 3 rats in the DLS (all with GRAB*rDA*3*m* and GRAB*ACh*), and 6 rats in the NAcc (all with GRAB*_DA_*_2_*_h_*). For behavioural analysis in Fig. 1 and Fig. 4, the 16 DMS rats were used, except in Fig. 4c. Among the 8 rats used for electrophysiology experiment, 3 rats were TH-Cre and 1 rat was Adora2a transgenic. No behavioural difference in task performance was observed between transgenic and wild-type rats.

Rats were water restricted to motivate them to perform the task. At the end of the day, rats were given 20 minutes of access to ad libitum water. Following behavioural sessions on Friday until mid-day Sunday, rats received ad libitum water. Animal use procedures were approved by the New York University Animal Welfare Committee (UAWC #2021-1120) and carried out in accordance with National Institute of Health standards.

### Behaviour training and task logic

Rats were trained in incremental stages of task complexity in a high-throughput behavioural facility using a computerized training protocol as described previously^28^.

LED illumination from the central nose port indicated that the animal could initiate a trial by poking its nose in that port (“CP On”). Upon trial initiation the center LED turned off. While in the central nose port, rats needed to maintain center fixation for a duration drawn uniformly from [0.8, 1.2] seconds. During the fixation period, a tone played from both speakers, the frequency of which indicated the volume of the offered water reward for that trial [1, 2, 4, 8, 16 kHz, indicating 5, 10, 20, 40, 80 µL rewards] (“Offer Cue”). Following the fixation period, one of the two side LEDs was illuminated, indicating that the reward might be delivered at that port (“Reward Port On”); the side was randomly chosen on each trial. This event also initiated a variable and unpredictable delay period, which was randomly drawn from an exponential distribution with mean = 2.5 seconds. The reward port LED remained illuminated for the duration of the delay period, and rats were freely behaving during this period. When reward was available, the reward port LED turned off (“Reward Cue”), and rats could collect the offered reward by nose poking in that port (“Reward Delivery”). The rat could also choose to terminate the trial at any time by nose poking in the opposite, un-illuminated side port (“Opt Out”), after which a new trial would immediately begin. On a proportion of trials (15-25%), the delay period would only end if the rat opted out (catch trials). If rats did not opt-out within 100 seconds on catch trials, the trial would terminate.

The trials were self-paced: after receiving their reward or opting out, rats were free to initiate another trial immediately. However, if rats terminated center fixation prematurely, they were penalized with a white noise sound and a time out penalty, typically lasting 2 seconds (“Violation”). The trial following a premature fixation break offered the rats the same amount of reward in order to dis-incentivize premature terminations for small volume offers.

We introduced semi-observable, hidden-states in the task by including uncued blocks of trials with varying reward statistics^74^: high and low blocks, which offered the highest three or lowest three rewards respectively, were interspersed with mixed blocks, which offered all volumes. There was a hierarchical structure to the blocks, such that high and low blocks alternated after mixed blocks (e.g., mixed-high-mixed-low, or mixed-low-mixed-high). Each session always started with a mixed block. Blocks transitioned after 40 successfully completed trials. Because rats prematurely broke fixation on a subset of trials, in practice, blocks varied in number of trials. Rats were considered to have achieved task proficiency when their time to opt-out exhibited linear sensitivity to the offered reward volume and time to opt-out for the 20 µL offer in low blocks was longer than in high blocks for at least 3 consecutive days. Rats that satisfied these criteria underwent surgeries specific to experiments. After recovering from surgeries, rats were retrained until they achieved the same pre-surgery performance which typically took one session. Rats were then habituated to being tethered for 2-3 days before proceeding with recordings and/or manipulations.

### Surgery

Rats were anaesthetized (isofluorane at 4% for induction and 1.5-2.5% for maintenance) and placed in a stereotaxic frame. Skull screws were inserted to secure implants before proceeding with surgeries specific to experiments.

#### Fibre photometry

For fibre photometry experiment with separate dopamine (N = 6 rats) and acetylcholine (N = 6 rats) recordings, rats were injected with 300-480 nL of either GRAB*_DA_*_2_*_h_* (AAV9-hSyn-GRAB DA2h, AddGene #140554) or GRAB*_ACh_* (AAV9-hSyn-ACh4.3, WZ Biosciences #YL10002) into the DMS (AP: −0.4 mm, ML: ±3.5 mm, DV: −4.6 mm or AP: 1.7 mm, ML: ±2.8 mm, DV: −4.3 mm from the skull surface at bregma at a 10^◦^ angle). To allow for brain motion correction, mCherry (AAV9-CB7-CI-mCherry-WPRE-RBG, AddGene #105544) was injected as a control fluorophore by mixing with the GRAB sensors at a ratio of 1.5:1-2.5:1 prior to injection. One rat injected with GRAB*_ACh_* did not receive mCherry and an isosbestic signal was used as the control.

For fibre photometry experiment involving simultaneous dopamine and acetylcholine recordings (N = 4 rats), rats were injected with a total of 1000-1080 nL of GRAB*_rDA_*_3*m*_ (pAAV9-hsyn-rDA3m, WZ Biosciences #YL002032-AV9) and GRAB*_ACh_* (AAV9-hSyn-ACh4.3, WZ Biosciences #YL10002) mixed at a ratio of 1:1-1:1.25.

All rats were then bilaterally implanted with 400 µM core, 0.5 NA optical fibres (Thorlabs) above the injection area. Craniotomies were sealed with Kwiksil and fibres were secured to the skull using dentin and Metabond. Rats were allowed to recover for a minimum of 7 days before behavioural training resumed.

#### Electrophysiology

The day before surgery, Neuropixels probe assembly was prepared by mounting the probe (Neuropixels probe 1.0) onto a custom-designed 3D printed piece and staining with DiI. Open Ephys built-in self tests were run on the probe assembly prior to surgery. Rats were implanted with a probe assembly in the DMS (AP: −0.24 mm, ML: ± 2.5 mm, DV: −4.4 mm from the skull surface at bregma; 6 rats in the left hemisphere, 2 rats in the right hemisphere). A ground wire was inserted in the cerebellum and fixed with Dentin. Craniotomies were covered with Kwik-Cast and gaps between the probe assembly and the skull were sealed with sterile paralube vet ointment. The probe was then fixed to the skull with Metabond and dental acrylic. Rats were allowed to recover for a minimum of 5 days before behavioural training resumed.

### Markerless pose tracking

Videos and synchronization pulses were acquired using methods described below for each experiment type. Deeplabcut software (v.2.2.3) was used for tracking the behaviour ports (right port, central port, and left port) and 6 body parts (right ear, nose, left ear, mid-point along the right torso, mid-point along the left torso, and base of the tail). Frames to be annotated and used as a training set were selected using k-means clustering. For each training cycle, 100-200 frames were extracted and annotated. The training was initialized with ResNet-50 and run for 500,000 iterations. Upon completion of training, the model-predicted positions of the features of interest were visually inspected. Outlier frames were then extracted manually and re-annotated. Frame annotation and training was repeated until no significant outliers existed, which typically took 4-5 training cycles with a total of 400-600 annotated frames.

The tracked features were smoothed in time with a moving median filter of 0.167 s. To obtain head speed and angle relative to the behaviour ports for analysis, the centroid of the head was calculated by averaging 3 body parts on the head (earR, earL, and nose). Extreme outliers in head speed and angle were removed by truncating the 99th percentile. Head speed and angle were aligned to task events by aligning the synchronization pulses with Bpod behaviour events.

### Fibre photometry experiment

#### Video and fibre photometry data acquisition

Video data were acquired using ceiling-mounted cameras to capture a top-down view of the arena (Doric USB3 behaviour camera, Sony IMX290, recorded with Doric Neuroscience Studio v6 software). To synchronize with neural recordings and Bpod task events, the camera was connected to a digital I/O channel of a fibre photometry console (FPC) and triggered at 30 Hz via a TTL pulse train. The remaining 3 digital I/O channels of the FPC were connected to the 3 behaviour ports to record TTL pulses when port LEDs turned on and off. An analog channel of the FPC was connected to low-autofluorescence patch cords to record fluorescence. Fluorescence from activity-dependent (GRAB*_DA_* and GRAB*_ACh_*) and activity-independent (isosbestic or mCherry) signals was acquired simultaneously via demodulation and downsampled on-the-fly by a factor of 25 to ∼481.9 Hz. Excitation wavelengths were 405 nm (isosbestic), 470 nm (GFP), and 560 nm (RFP). For three-channel acquisition used in simultaneous dopamine and acetylcholine recordings, the RFP excitation wavelength was 568 nm. The photometry rig included a rotary fluorescence minicube (RFMC6 IE(400-410) E1(460-490) F1(500-540) E2(550-570) F2(580-680) S). The power at the tip of the fiber optic was 20-30 µW for each excitation channel.

#### Fibre photometry data preprocessing

The recorded demodulated fluorescence was corrected for photobleaching and motion using Two-channel Motion Artifact Correction (TMAC)^33^ with mCherry or isosbestic signal as the activity-independent channel. TMAC uses a Bayesian inference-based algorithm to infer motion artifacts from the control channel (i.e., an activity-independent indicator). For the cohort of rats where dopamine and acetylcholine were recorded separately, 9 out of 10 rats were injected with mCherry. For the cohort injected with GRAB*_ACh_* and GRAB*_rDA_*_3_*_m_* for simultaneous recording, isosbestic signal of GRAB*_ACh_* was used as the control channel. The excitation wavelength we used for the isosbestic point is based on previously characterized properties of the GRAB*_ACh_*_3.0_ sensor, and what has been used as the isosbestic point for this sensor in past studies^75^. The motion-corrected signal was z-scored and then aligned to task events using synchronization TTL pulses from the behaviour ports. It is possible that the GRAB*_ACh_*_4.3_ sensor we used has a slightly shifted isosbestic point from the GRAB*_ACh_*_3.0_ sensor^75^. However, the fact that results were qualitatively the same regardless of motion correction with mCherry versus the isosbestic point shows that this did not impact the results or conclusions of our study (Extended Data Fig. 7, 8).

### Electrophysiology experiment

#### Neuropixels data acquisition

Spike data was acquired from 384 channels at 30 kHz using Neuropix-PXI hardware and OpenEphys. To synchronize with behaviour, task event TTL pulses were recorded simultaneously using NI-DAQmx.

#### Spike sorting

Spike data were preprocessed using Kilosort 2.5. After preprocessing, clusters that were identified from Kilosort as single-units were manually inspected using the Phy2 Python package. Clusters with unreliable waveform shape, autocorrelogram, or spike amplitude timecourse were manually classified as multi-unit or noise and removed from further analysis. For the identified single-units, spike data were aligned to behavioural events and binned into 5 ms bins using custom Matlab codes.

### Event-aligned photometry signals

To align photometry signals to task events, motion-corrected signals were z-scored within each session before pooling sessions for each rat. Event-aligned responses were then averaged across rats. For analyses split by reward delay quartiles, trials with reward delays shorter than 0.75 s were excluded to avoid bleed-over responses from Reward Port On event. When baseline correction was applied, the minimum value of the condition-averaged signal (e.g., different offer volumes, delay quartiles, reward port side) during the baseline window (0.1 to 0 s for all events except opt-out; −0.3 to 0 s for opt-out) was subtracted from all trials corresponding to that condition.

### Quantification of task variable encoding

To quantify dopamine encoding of task variables for Extended Data Fig. 3e and 5d, we computed the discriminability index (*d*^′^) of event-aligned dopamine responses, conditioned on the offered reward volume, reward delay, and reward port side. Since reward port side is a binary variable (contralateral vs. ipsilateral), we grouped the other task variables into two categories for consistency: for reward volume, we compared 5 and 10 µL with 40 and 80 µL offer trials; for reward delay, we compared the bottom 50% vs. top 50% of the reward delay distribution. For each rat, we first computed the average z-scored event-aligned signal for each condition. Then, the discriminability index was calculated as: 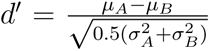 where *µ* and *σ* are the mean and standard deviation, respectively, of the distribution across sessions.

### Behavioural model

We modeled the trial initiation times as a linearly decreasing function of the value of the environment, which was updated according to the following recursive equation: *V_t_*_+1_ = *V_t_* + *αRPE*. The RPE was defined as *R_t_* −*V_t_* for the volume RPE model and *Delay_t_* −*V_t_* for the delay RPE model. The learning rate *α* ∈ [0, 1] governed the rate at which value estimates were updated across trials. Trial statistics from the same rats used in behavioral analyses were used to simulate the model. Best-fit parameters for the learning rate *α* and the scaling factor *β*, which determined the relationship between value and trial initiation time, were obtained by simulating a grid of parameter combinations and selecting those that qualitatively best captured rats’ behaviour. The final parameter pairs used were (*α, β*) = (0.5, 0.48) for the volume RPE model and (0.46, 0.54) for the delay RPE model.

### Trial initiation vs dopamine AUC analysis

To relate dopamine at the offer cue to trial-by-trial changes in trial initiation time (Fig. 3e), we restricted the analysis to post-violation trials, as prior work showed that reward volume modulation of initiation time is driven primarily by these trials[28]. Specifically, we analyzed how dopamine on a violation trial N related to changes in trial initiation time between trials N and N+1.

For each rat, we removed outliers in initiation times by excluding the bottom and top 10% of values pooled across all sessions. We then computed the change in trial initiation times as: -Δ initiation = -(initiation time*_N_*_+1_ - initiation time*_N_*). A positive value of -Δ initiation means that the rat initiated the subsequent trial faster. -Δ initiation values were then split into 6 bins: the negative and positive values were each divided into terciles, resulting in the first 3 bins for trials with subsequently slower initiation and the last 3 bins for subsequently faster initiation.

To compute dopamine AUC, z-scored photometry signal was aligned to the offer cue and baseline corrected using the same method described in *Event-aligned photometry signal* (see Methods). AUC was calculated from 0 to 0.5 s following offer cue onset on each trial. Trials with extreme AUC values (bottom and top 0.1%) were excluded. For each rat, the average AUC was computed within each of the 6 initiation bins and then normalized by z-scoring across bins before pooling across rats.

For the reward cue analysis (Fig. 3h), the procedure was identical except that dopamine AUC was aligned to the reward cue, and trial initiation changes were calculated from a given reward cue trial N to trial N+1.

### Trial-by-trial spiking analysis

To account for biases in statistical test due to imbalanced dataset across task events, we performed random upsampling of the minority dataset. If the ratio of the number of data points between the majority and the minority datasets exceeded 2.5, the minority dataset was upsampled via random sampling with gaussian noise (sigma = 1). Only sessions that had at least 20 trials were included for analysis.

### Logistic regression model

To compare encoding of movement vigour in DMS spikes, we trained a logistic regression decoder. For each session, the predictor tensor matrix (Trial x Neuron x Time) was built by concatenating spikes from 0 to 0.5 seconds after the onset of reward port assignment. For each trial in a given session, it was classified as fast/slow if it were in the bottom/top quartile of the reaction time distribution. Only sessions with at least 5 simultaneously recorded DMS units were included for analysis. We used the Matlab function fitclinear to train the decoder to predict 1 if a given trial is fast, or 0 if slow. To cross-validate, we implemented 10-fold cross-validation with ridge regression. The regularization strength was determined by training the decoder sequentially on a set of 11 logarithmically-spaced regularization strengths ranging from 10^−4^ to 10^−0.2^ and selecting the value that minimizes classification error.

### Histology

Following completion of each experiment, rats were deeply anesthetized and transcardially perfused with phosphate buffer saline (PBS) and formalin for histological verification. Brains were post-fixed with formalin for 3 days and sectioned at 50 *µ*m coronal slices on a Leica VT1000 vibratome. For electrophysiology experiment, sections were imaged with an Olympus VS120 Virtual Slide Microscope to verify probe location. For fibre photometry experiment, sections were stained to enhance GFP and imaged to verify virus expression and fibre implant location.

#### Immunohistochemistry

For brain tissues acquired for fibre photometry experiment, sections were treated with 1% hydrogen peroxide in 0.01 M PBS for 30 min, washed with 0.01 M PBS, and blocked in 0.01 M PBS containing 0.05% sodium azide and 1% bovine serum albumin (PBS-BSA-Azide) for 30 minutes prior to treating with primary antibodies. GFP was amplified with Rabbit anti-GFP (Thermo Fisher Scientific A11122, 1:2,000, lot #2083201) as primary antibody and Goat anti-Rabbit AlexaFluor 488 IgG (Thermo Fisher Scientific A11008, 1:200, lot #2147635) as secondary antibody. Primary and secondary antibodies were made in PBS-BSA-Azide and primary antibodies additionally contained 0.2% triton-x. Sections were incubated in primary antibody overnight at room temperature, washed with PBS for 15 min, and treated with secondary antibody for 1 hour in dark. After washing with PBS for 15 min, sections were mounted on glass slides using Prolong Gold (Vector Laboratories).

## Acknowledgments

We thank Tanya Sippy, Cristina Savin, Robert Froemke, Richard Tsien, Mike Tadross, and members of the Constantinople lab for feedback. We thank research technicians in the Constantinople lab for animal training. We thank Juan Mena-Segovia for providing transgenic animals for revision experiments.

## Author Contributions

H.J.J. and C.M.C. designed the study. H.J.J., R.M.W., and C.E.G. collected data and H.J.J. analyzed data. C.M.C. advised on the data analysis. H.J.J. and C.M.C. wrote the paper. C.M.C. supervised the project.

