## Supplemental Figures for "Acetylcholine demixes heterogeneous dopamine signals for learning and moving"

### Supplementary materials

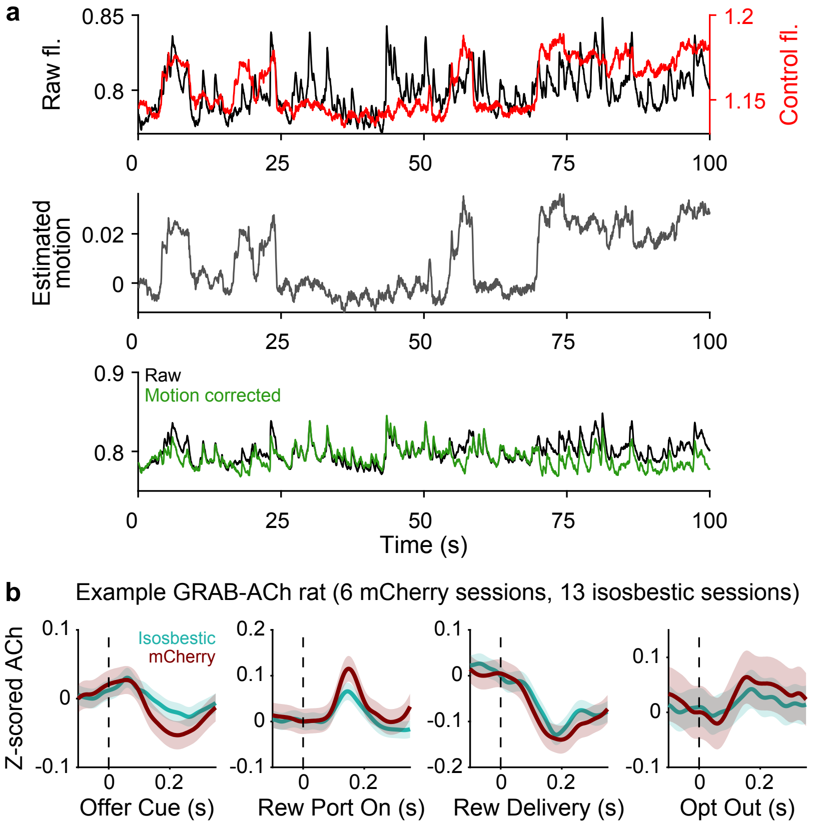


Extended Data Fig. 1: **Motion correction using TMAC produces similar event-aligned responses for both mCherry and isosbestic control. a.** Top, Example raw fluorescence (black) overlaid with simultaneously recorded mCherry fluorescence (red). Middle, Estimated motion calculated with TMAC (grey). Bottom, Raw fluorescence overlaid with TMAC motion corrected fluorescence (green). **b.** Average motion-corrected acetylcholine signal using isosbestic (green, N = 13 sessions) or mCherry (red, N = 6 sessions) as the activity-independent channel for an example rat.

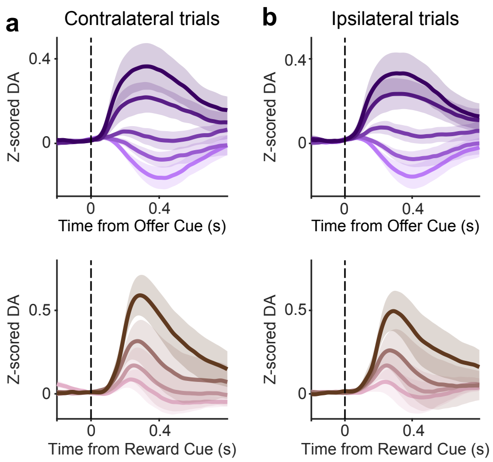


Extended Data Fig. 2: **Dopamine release is not side selective at RPE-associated events. a.** Z-scored dopamine release split by offered reward volume in mixed blocks at the time of the offer cue (top) and by reward delay quartile at reward cue (bottom) on contralateral trials. **b.** Same as in **a** but on ipsilateral trials. Across panels, N = 10 rats, mean ± s.e.m..

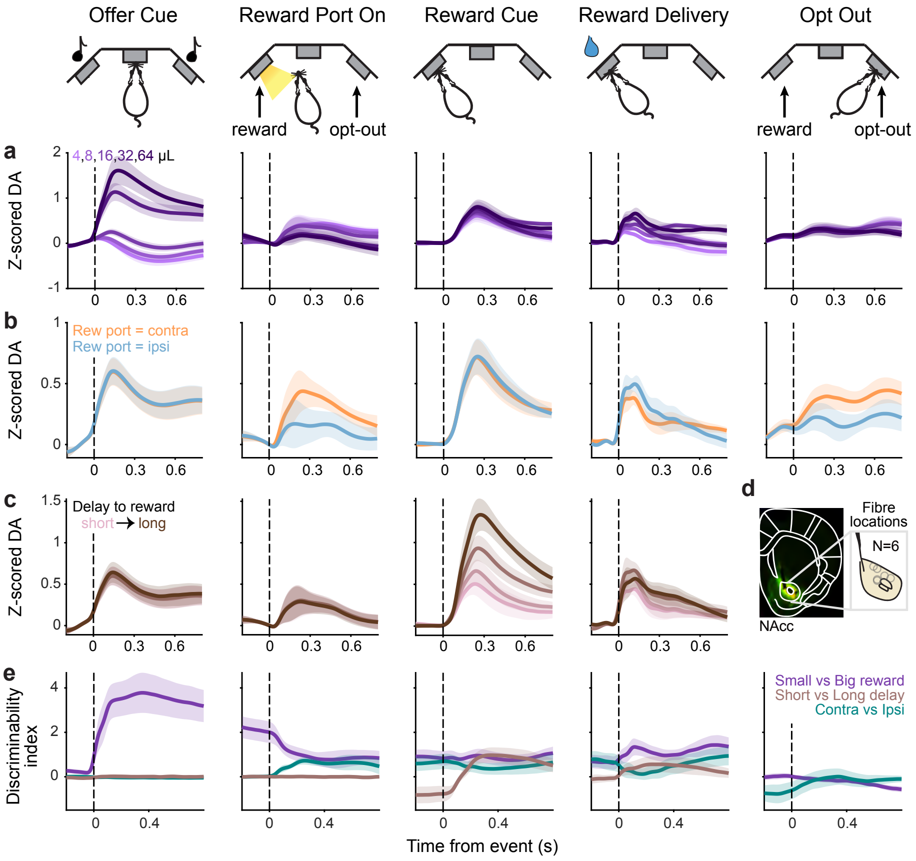


Extended Data Fig. 3: **Dopamine dynamics in the NAcc. a.** Average event-aligned dopamine release in the NAcc split by offered reward volume in mixed blocks. These data were collected from female rats, so they were offered slighty smaller water volumes compared to males, to obtain sufficient numbers of behavioural trials[28,30] (4-64 $\mu$L versus 5-80 $\mu$L for males). **b.** Average event-aligned dopamine release in the NAcc split by reward delay quartiles across all blocks. **c.** Average event-aligned dopamine release in the NAcc split by contralateral and ipsilateral trials. **a.-c.,** Dopamine signals are z-scored and baseline-corrected before pooling across rats. **d.** Representative histology image of viral expression of GRAB$_{DA2h}$ and mCherry and recovered fibre placements. **e.** Average discriminability index of different task variables (purple: reward volume, brown: reward delay, green: contra/ipsi) aligned to all task events. N = 6 rats, mean ± s.e.m. across panels.

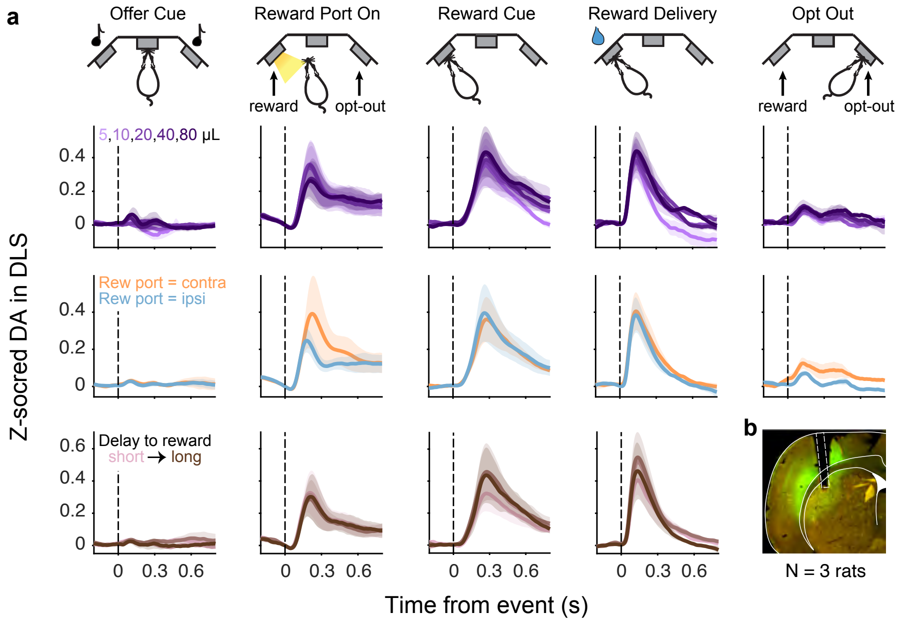


Extended Data Fig. 4: **Dopamine dynamics in the DLS. a.** Event-aligned dopamine release in the DLS for different reward offers during mixed blocks (top), reward port location (middle), and delay to reward quartiles (bottom), averaged across rats (N = 3 rats, mean ± s.e.m.). These rats were recorded with medium-affinity red-shifted GRAB sensor (rDA3m). Dopamine signals are z-scored and baseline corrected before pooling across rats. **b.** Representative histology image of viral expression of $GRAB_{rDA3m}$ and $GRAB_{ACh}$ and fibre track in the DLS.

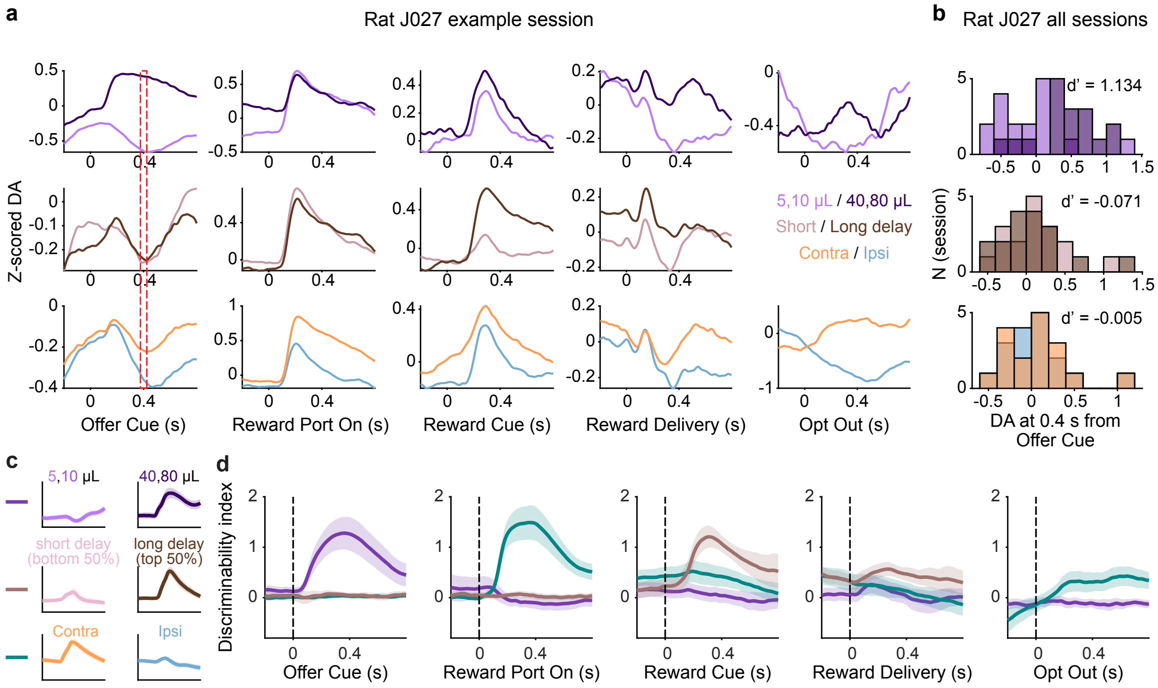


Extended Data Fig. 5: **Discriminability index quantifies encoding of task variables at different events. a.** Top, Average z-scored dopamine release on small (5, 10 µL) versus large (40, 80 µL) reward offer trials aligned to all task events in an example session. Middle, Average z-scored dopamine release on short (bottom 50% of reward delay distribution) vs long (top 50%) reward delay trials. Bottom, Average z-scored dopamine release on contralateral versus ipsilateral trials. Across panels, the red vertical line indicates the time point the histograms in **b** were generated from (t = 0.4 s from the offer cue onset). **b.** Distribution of z-scored dopamine at 0.4 s from the offer cue onset (red vertical line in **a**) across 17 sessions from an example rat. **c.** Schematic of how task variables are binarized to compute discriminability index. **d.** Average discriminability index of different task variables (purple: reward volume, brown: reward delay, green: contra/ipsi) aligned to all task events (N = 10 rats, mean ± s.e.m.).

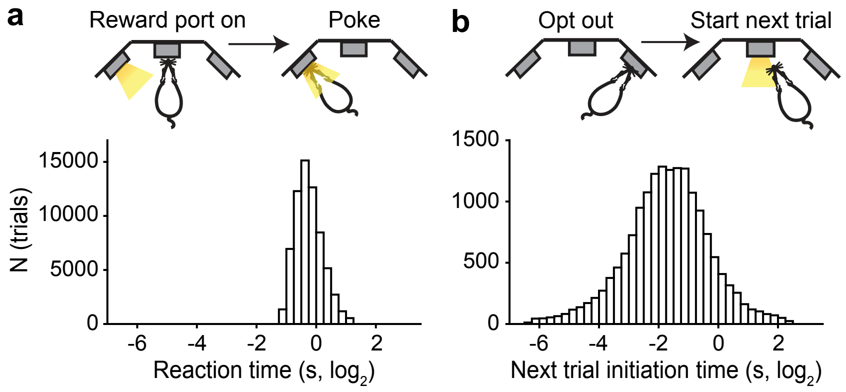


Extended Data Fig. 6: **Behavioural variability is higher at opt-out compared to reward port assignment. a.** Histogram of rats' reaction time to the side LED lighting up to indicate the side of the reward port. **b.** Histogram of rats' time to initiate trial following opt-outs. N = 16 rats across panels.

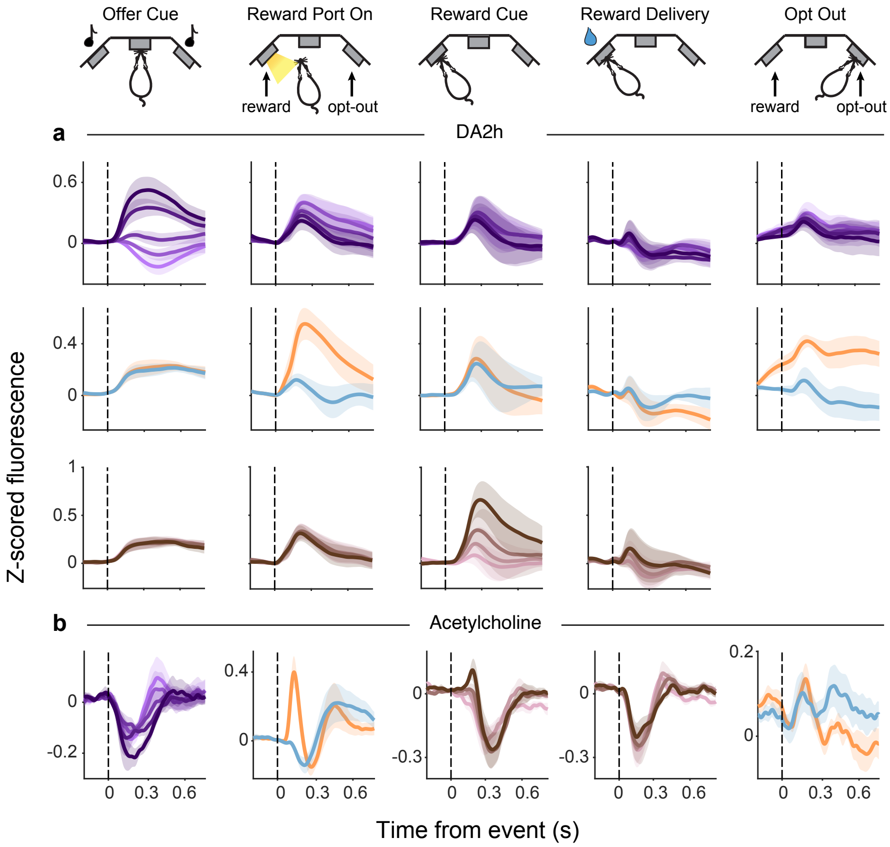


Extended Data Fig. 7: **Acetylcholine and high-affinity dopamine sensor dynamics in the DMS show characteristic RPE and contralateral selectivity when recorded in separate animals.** **a.** Event-aligned z-scored dopamine signals split by reward offer volume (top, light to dark purple for small to large volumes), reward port location (middle, orange for contralateral and blue for ipsilateral), and delay to reward quartile (bottom, pink to brown for shortest to longest delay quartile bin), averaged across rats (N = 6 rats, mean ± s.e.m.). **b.** Same as in **a** but for acetylcholine (N = 6 rats, mean ± s.e.m.). Across panels, signals are baseline corrected (Methods).

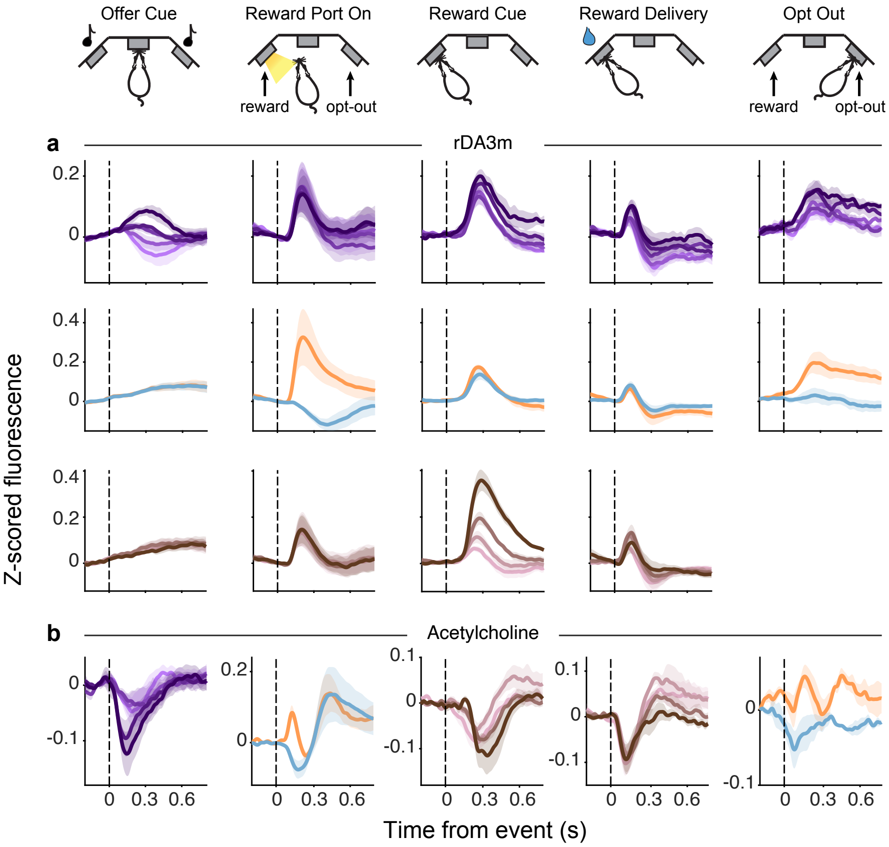


Extended Data Fig. 8: **Acetylcholine and medium-affinity dopamine sensor dynamics show characteristic RPE and contralateral selectivity when recorded simultaneously. a.** Event-aligned z-scored dopamine signals split by reward offer volume (top, light to dark purple for small to large volumes), reward port location (middle, orange for contralateral and blue for ipsilateral), and delay to reward quartile (bottom, pink to brown for shortest to longest delay quartile bin), averaged across rats. **b.** Same as in **a** but for acetylcholine. Across panels, N = 4 rats, mean ± s.e.m., baseline corrected (Methods).

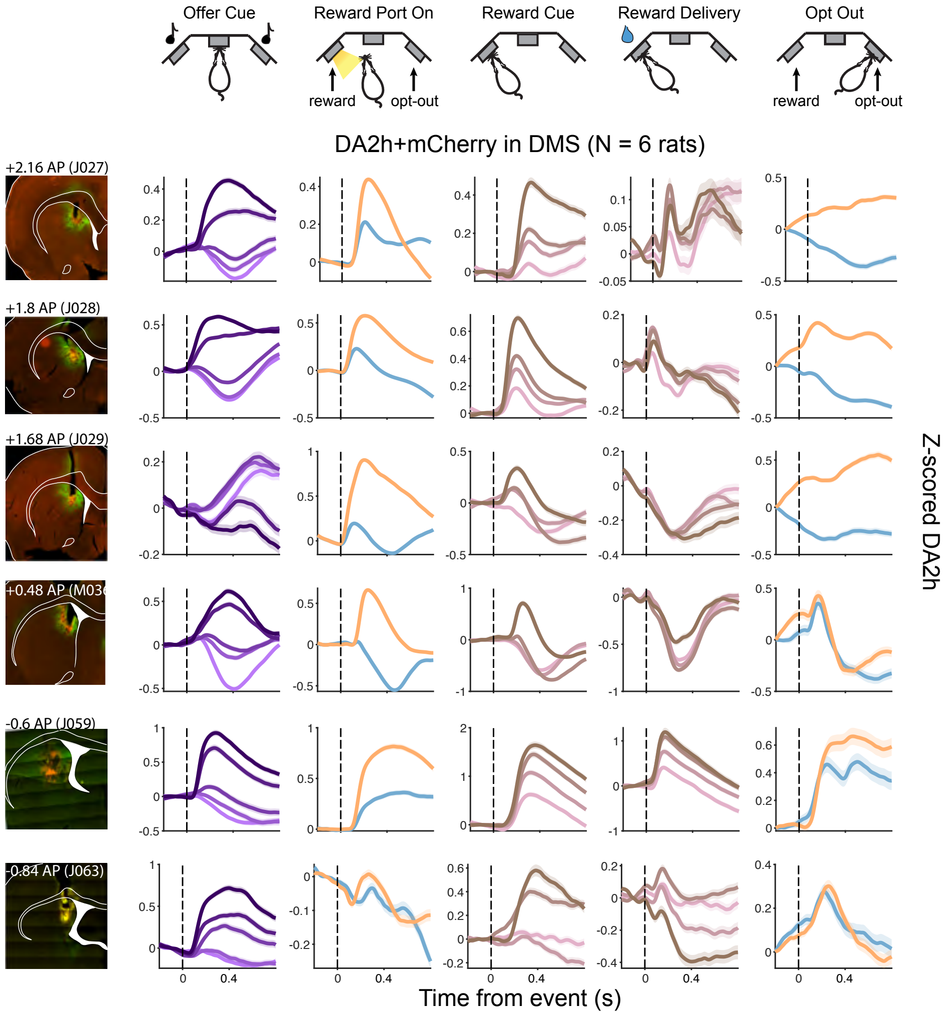


Extended Data Fig. 9: **Characteristic dopamine RPE responses and contralateral selectivity are present across the DMS.** Each row shows histology and event-aligned dopamine release for individual rats injected with green-fluorescent DA2h and mCherry (N = 6 rats). Event-aligned signals are z-scored and baseline corrected.

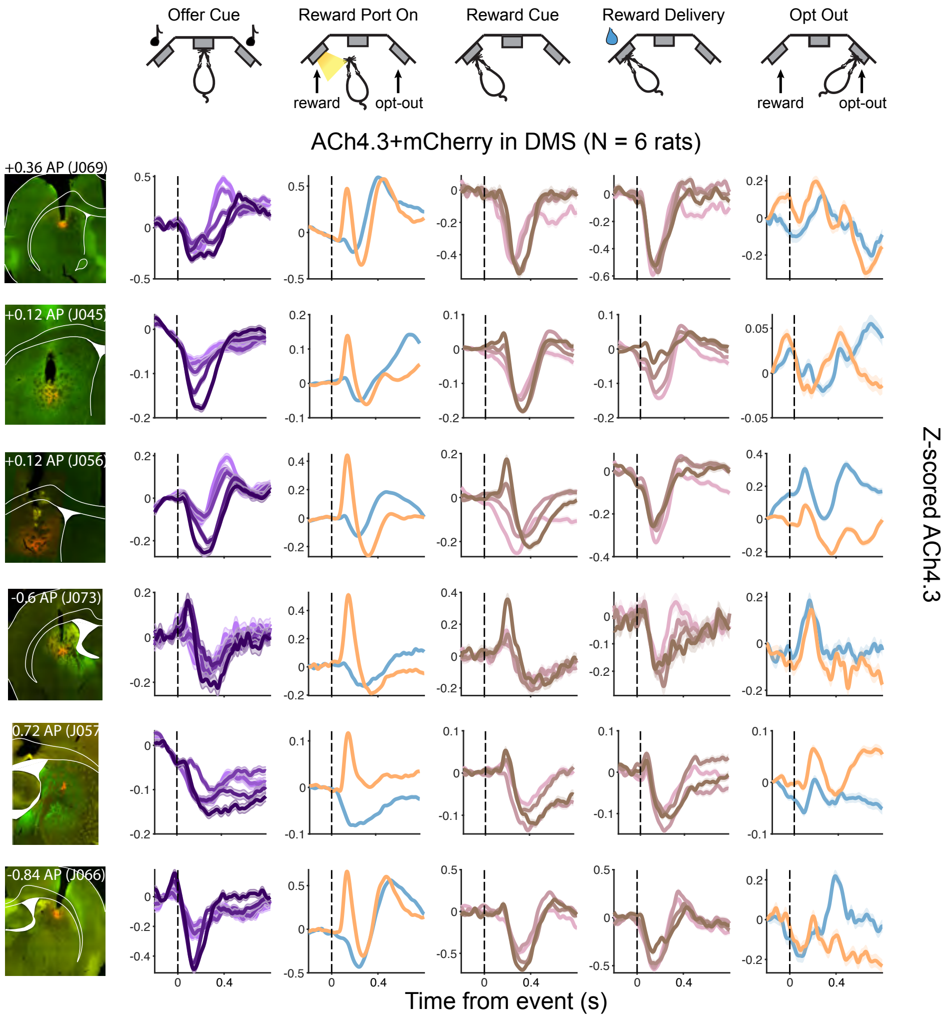


Extended Data Fig. 10: **Characteristic cholinergic dips and bursts at RPE- and movement-associated events are present across the DMS.** Each row shows histology and event-aligned acetylcholine release for individual rats injected with green-fluorescent ACh4.3 and mCherry (N = 6 rats). Event-aligned signals are z-scored and baseline corrected.

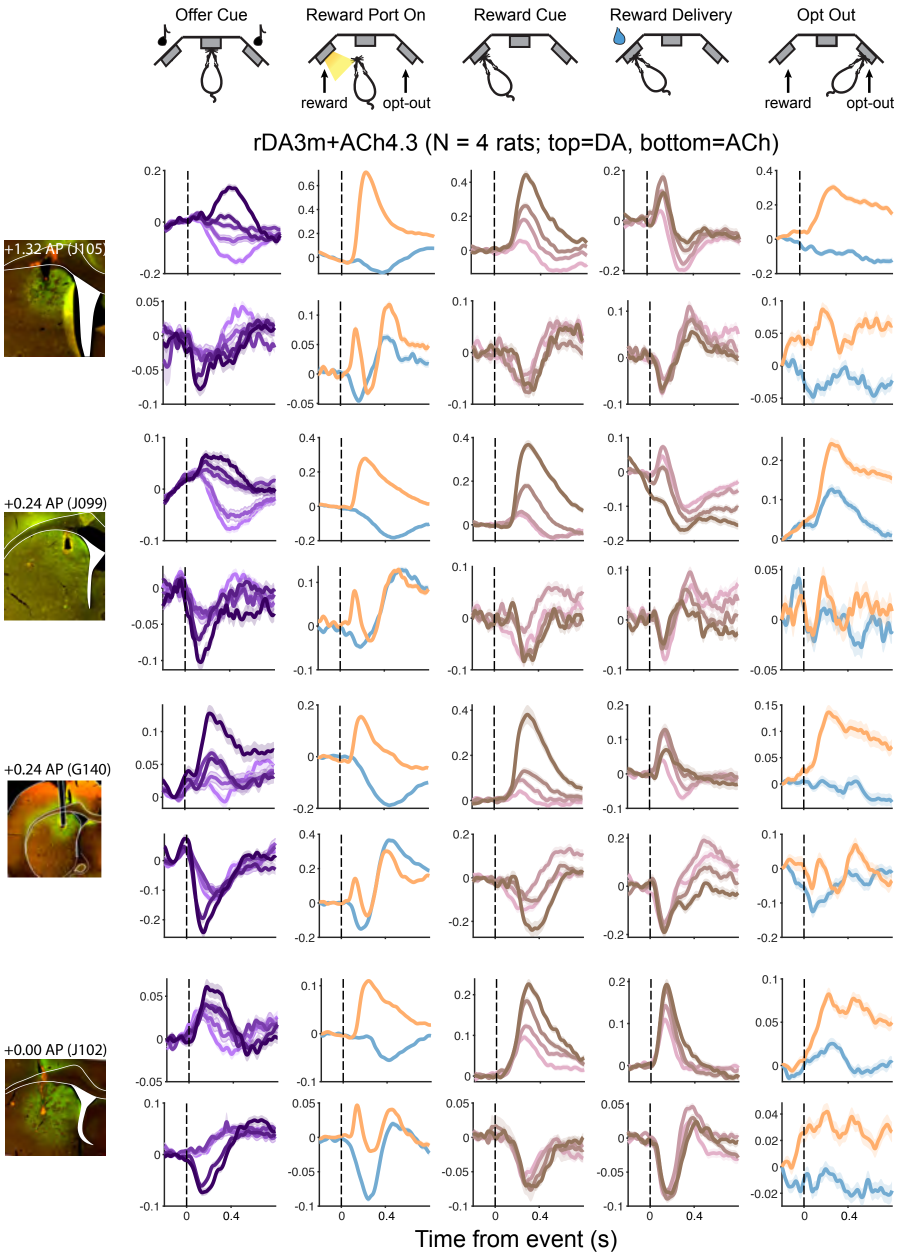


Extended Data Fig. 11: **Simultaneous measurement of dopamine and acetylcholine release reveals characteristic dopamine and acetylcholine dynamics at RPE- and movement-associated events across the DMS.** Event-aligned dopamine (top) and acetylcholine (bottom) responses in rats injected with red-shifted rDA3m and green-fluorescent ACh4.3 (N = 4 rats). Event-aligned signals are z-scored and baseline corrected.

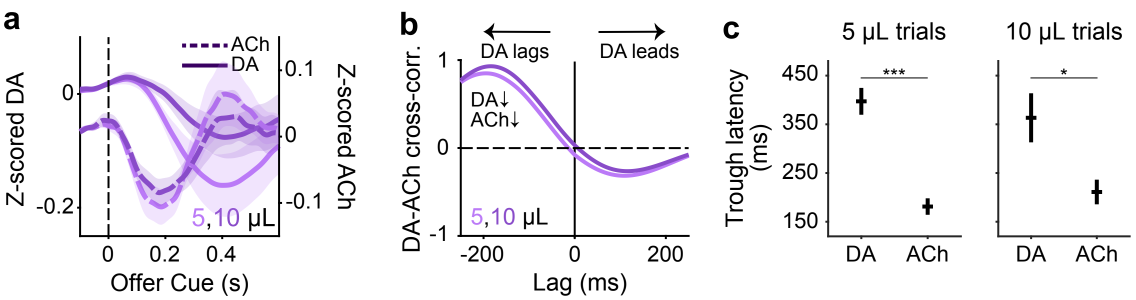


Extended Data Fig. 12: **Acetylcholine dips slightly lead dips in dopamine at the offer cue.** **a.** Z-scored dopamine (solid, N = 10 rats) and acetylcholine (dashed, N = 10 rats) signals for small reward offer trials in mixed blocks aligned to the onset of offer cue, averaged across rats. **b.** Cross-correlation between average dopamine and acetylcholine signals around offer cue (-0.1-0.5 s) for small reward offer trials. Negative lag indicates dopamine following acetylcholine. **c.** Latency to trough in dopamine and acetylcholine release after the onset of offer cue on 5 μL (left) and 10 μL (right) offer trials in mixed block, averaged across rats (N = 10 DA rats, N = 10 ACh rats, mean ± s.e.m.; two-sided Wilcoxon rank sum test: 5 μL trials, $p < 0.001$; 10 μL trials, $p = 0.0140$).

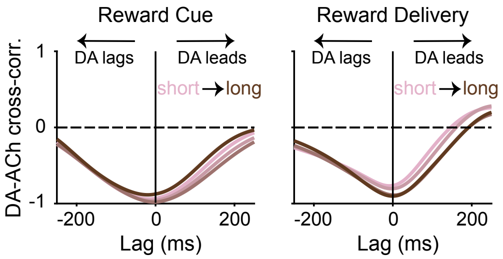


Extended Data Fig. 13: **Acetylcholine and dopamine dynamics in the DLS are antiphase at the time of reward cue and delivery.** Cross-correlation of rat-averaged dopamine and acetylcholine signals (-0.1-0.5 s) at the reward cue (left) and reward delivery (right) on trials with different reward delay quartiles. N = 3 rats.

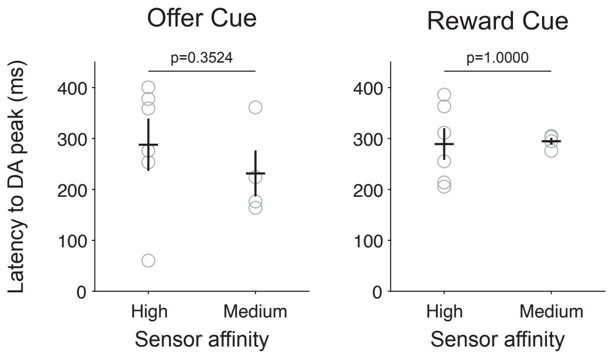


Extended Data Fig. 14: **Latency to peak in dopamine at reward-associated events does not depend on sensor affinity.** Latency to peak in dopamine at the time of the offer cue (left) and reward cue (right) by sensor affinity, averaged across rats (N = 10 rats for high-affinity, N = 4 rats for medium-affinity, mean ± s.e.m.; two-sided Wilcoxon rank-sum test: Offer Cue, p = 0.3524; Reward Cue, p = 1.0000). Grey circles indicate individual rats.

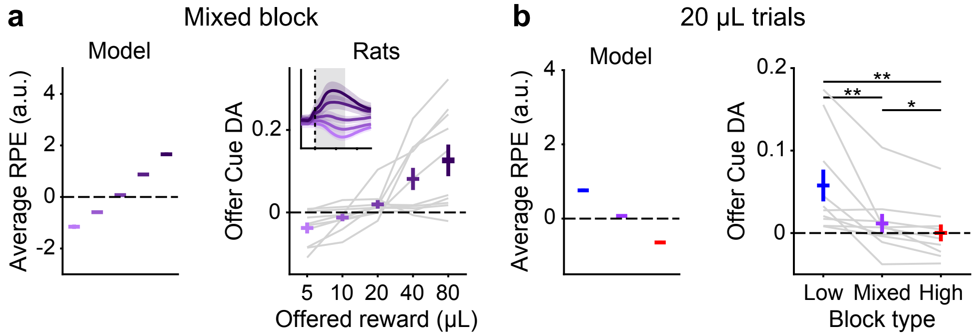


Extended Data Fig. 15: **Reward volume RPE model captures dopamine responses at the offer cue. a.** Left, Average model predicted RPE by offered reward volume in mixed blocks. Right, Dopamine AUC at the offer cue (0-0.5 s) in mixed blocks (N = 10 rats). Grey lines show individual rats. Mean ± s.e.m. across panels. **b.** Same as in **a** but on 20 μL offer trials by block type (N = 10 rats, mean ± s.e.m., two-sided Wilcoxon signed-rank: low vs mixed: p = 0.0039; low vs high: p = 0.0020; mixed vs high: p = 0.0273).

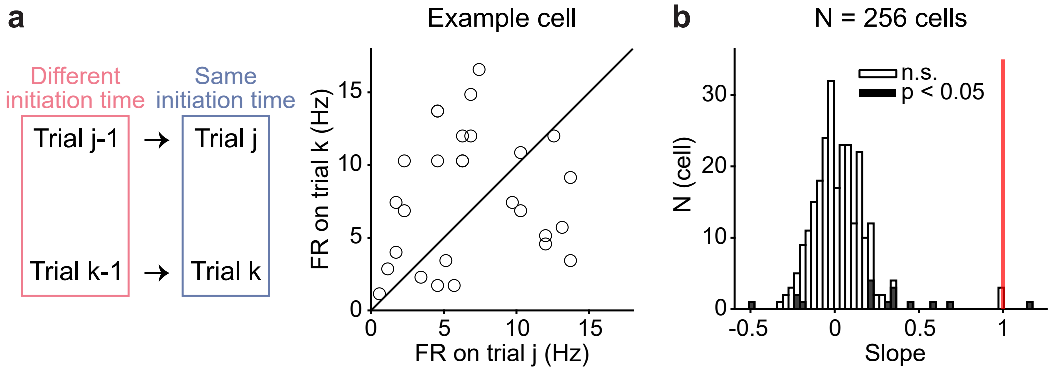


Extended Data Fig. 16: **Trial initiation speed on current trial alone does not explain DMS firing rate. a.** Left, Trials j and k have similar trial initiation times (difference $< 0.01$ s) but different initiation times on the previous trial (difference $> 2$ s). Right, Average firing rate at offer cue (0-0.3 s) on trials j and k for an example cell. Solid line is the unity line. **b.** Histogram of best-fit line slopes for all cells. Red vertical line indicates slope of 1.

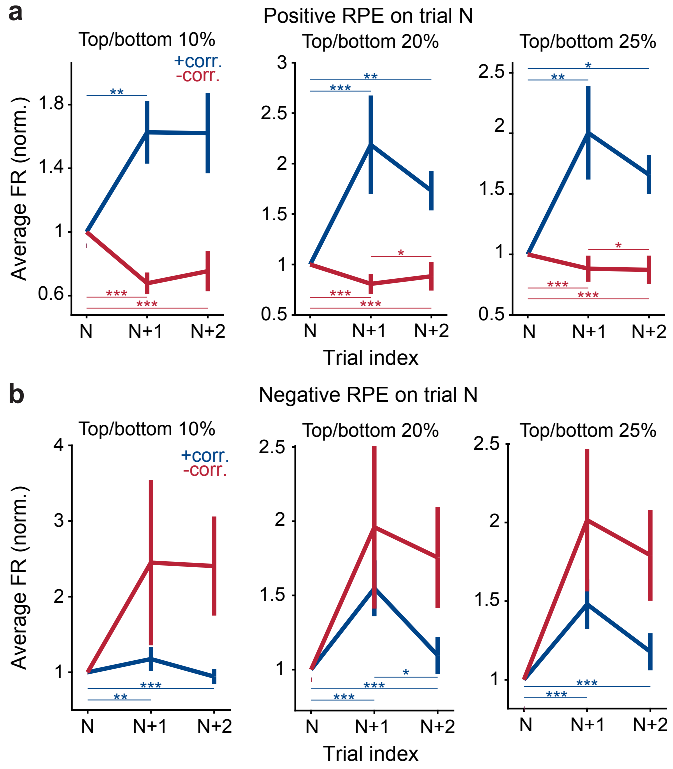


Extended Data Fig. 17: **RPE produces lasting changes in task-evoked firing rates for a range of RPE thresholds. a.** Change in average z-scored firing rate at offer cue (0-0.3 s) on 3 consecutive trials where a big positive RPE on trial N is followed by a negligible RPE on trial N+1. Different panels are for different thresholds used to select RPE on trial N. Two-sided Wilcoxon signed-rank: Left, 10% threshold: +corr. cells, N = 100 trial sequences, trial N vs N+1, p = 0.0064; trial N vs N+2, p = 0.0620; trial N+1 vs N+2, p = 0.4269; -corr. cells, N = 115 trial sequences, trial N vs N+1, $p < 0.001$; trial N vs N+2, $p < 0.001$; trial N+1 vs N+2, p = 0.5312. Middle, 20% threshold: +corr. cells, N = 191 trial sequences, trial N vs N+1, $p < 0.001$; trial N vs N+2, p = 0.0395; trial N+1 vs N+2, p = 0.1138; - corr. cells, N = 223 trial sequences, trial N vs N+1, $p < 0.001$; trial N vs N+2, $p < 0.001$; trial N+1 vs N+2, p = 0.0182. Right, 25% threshold: +corr. cells, N = 245 trial sequences, trial N vs trial N+1, p = 0.0025; trial N vs trial N+2, p = 0.0434; trial N+1 vs trial N+2, p = 0.3320; -corr. cells, N = 280 trial sequences, trial N vs trial N+1, $p < 0.001$; trial N vs trial N+2, $p < 0.001$; trial N+1 vs trial N+2, p = 0.0339. **b.** Same as in **a** but for a big negative RPE on trial N. Two-sided Wilcoxon signed-rank: Left, +corr. cells, N = 119 trial sequences, trial N vs trial N+1, p = 0.0010, trial N vs trial N+2, $p < 0.001$; Middle, +corr. cells, N = 240 trial sequences, trial N vs trial N+1, p = 0.0005; trial N vs trial N+2, $p < 0.001$; trial N+1 vs trial N+2, p = 0.0108; Right, +corr. cells, N = 299 trial sequences, trial N vs trial N+1, $p < 0.001$; trial N vs trial N+2, $p < 0.001$. Across panels, *$p < 0.05$, **$p < 0.01$, ***$p < 0.001$.

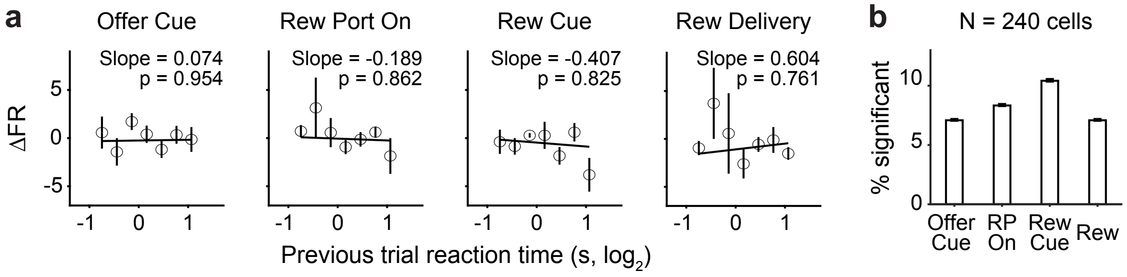


Extended Data Fig. 18: **Speed of contralateral movement does not modulate DMS firing rate on subsequent trial. a.** Trial-by-trial changes in average firing rate surrounding each task event (0-0.3 s) versus putative DA at Reward Port On (i.e., vigour of contralateral movement) for an example cell. Solid lines show the best-fit line (t-test on the slope: Offer Cue, p = 0.954; Reward Port On, p = 0.862; Reward Cue, p = 0.825; Rew Delivery = 0.761). Trials are binned in 7 linearly spaced intervals for visualization (mean ± s.e.m.). **b.** Fraction of cells with significant slopes at different task events. Error bars are binomial confidence intervals.

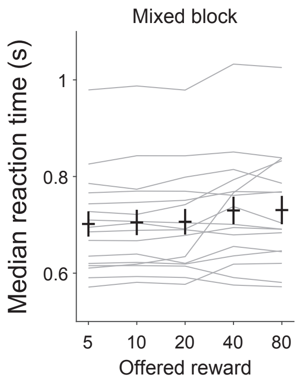


Extended Data Fig. 19: **Vigour of orienting to the reward port upon illumination does not correlate with the reward offer volume.** Median reaction time to the reward port LED illumination at the reward port assignment in mixed blocks for different reward offers, averaged across rats (N = 16 rats, mean ± s.e.m.; linear mixed-effects model fit on medain reaction time vs reward volume: slope = 0.0082, $p < 0.001$). Grey lines indicate individual rats. The lack of reward-dependent modulation of movement speed is consistent with this orienting movement reflecting a prepotent, reflexive response to a salient stimulus (the LED turning on).

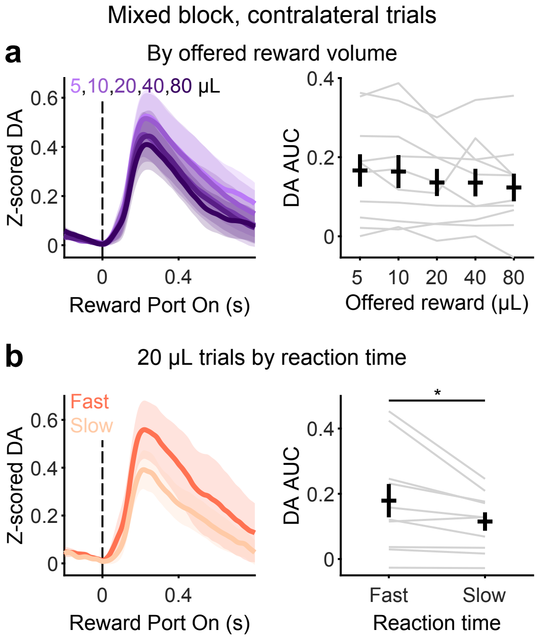


Extended Data Fig. 20: **Dopamine release at reward port assignment is not modulated by value. a.** Dopamine release (left) and AUC (right) for varying reward offer volumes on contralateral trials during mixed blocks, averaged across rats (N = 10 rats, mean ± s.e.m.). Grey lines indicate individual rats. **b.** Dopamine release (left) and AUC (right) for 20 µL contralateral trials during mixed blocks when the side LED orienting response is in the fastest versus slowest quartile (N = 10 rats, mean ± s.e.m.; two-sided Wilcoxon signed-rank: $p = 0.0137$). Grey lines indicate individual rats.
